# Impact of natural transformation on the acquisition of novel genes in bacteria

**DOI:** 10.1101/2025.01.24.634696

**Authors:** Fanny Mazzamurro, Marie Touchon, Xavier Charpentier, Eduardo PC Rocha

## Abstract

Natural transformation is the only process of gene exchange under the exclusive control of the recipient bacteria. It has often been considered as a source of novel genes but quantitative assessments of this claim are lacking. To investigate the potential role of natural transformation in gene acquisition, we analysed a large collection of genomes of *Acinetobacter baumannii* (Ab) and *Legionella pneumophila* (Lp) for which transformation rates were experimentally determined. Natural transformation rates are weakly correlated with genome size. But they are negatively associated with gene flow in both species. This might result from a negative balance between transformation’s ability to cure the chromosome from mobile genetic elements (MGEs), resulting in gene loss, and its facilitation of gene acquisition. By focusing on the latter, we found that transformation was significantly associated with small gene acquisition events while MGEs-driven gene acquisition tend to be associated with larger ones. Events of gene gain by transformation were spread more evenly in the chromosome than MGEs encoding the ability to integrate autonomously. We estimated the contribution of natural transformation to gene gains by comparing recombination-driven gene acquisition rates between transformable and non-transformable strains. Natural transformation may have caused the acquisition of up to 6.4% (Ab) and 1.1% (Lp) of the novel genes. This low contribution of natural transformation to the acquisition of novel genes implies that most novel genes must have been acquired by other means. Interestingly, the ones potentially acquired by transformation include almost 15% of the recently acquired antibiotic resistance genes in *A. baumannii*. Hence, natural transformation may drive the acquisition of relatively few novel genes but these may have a high fitness impact.

## Introduction

Horizontal gene transfer (HGT) accelerates the evolution of bacterial genomes by three main mechanisms: DNA conjugation, DNA transfer in viral particles (phages), and natural transformation. Among these three, only natural transformation is under the direct control of the recipient bacteria (1). During transformation, bacteria uptake exogenous DNA from their environment and integrate it in their chromosome by homologous recombination if it exists high sequence similarity between exogenous and native DNA (2). Recombination can result in a simple allelic exchange between homologous regions. It has been shown that naturally transformable bacteria tend to have higher rates of recombination leading to such exchanges in core genes (3). Yet, if there are two distinct regions of homology in the DNA sequence, recombination may result in the acquisition of novel genes from the exogenous DNA and cause gene deletions in the native chromosomal region (Figure 1). The exact outcome will then depend on the intervening sequence between the two homologous regions in the chromosome and in the incoming DNA (2). This process may be frequent, since it was previously shown that core genes next to recently acquired DNA show higher rates of recombination than the others (3,4). Furthermore, the potential of transformation to result in the acquisition of novel genes is known for almost a century. Natural transformation was discovered in 1928 by Griffith in *Streptoccocus pneumoniae* while observing conversions between virulent and non-virulent strains (5). The reported transforming principle was later demonstrated to be DNA (6) encoding the complete capsule locus (5,7). Since then, the core components of the machinery necessary for DNA uptake by natural transformation were uncovered (2,8,9). A wide variety of bacterial species are competent and naturally transformable (10), among which numerous pathogens.

**Figure 1.**
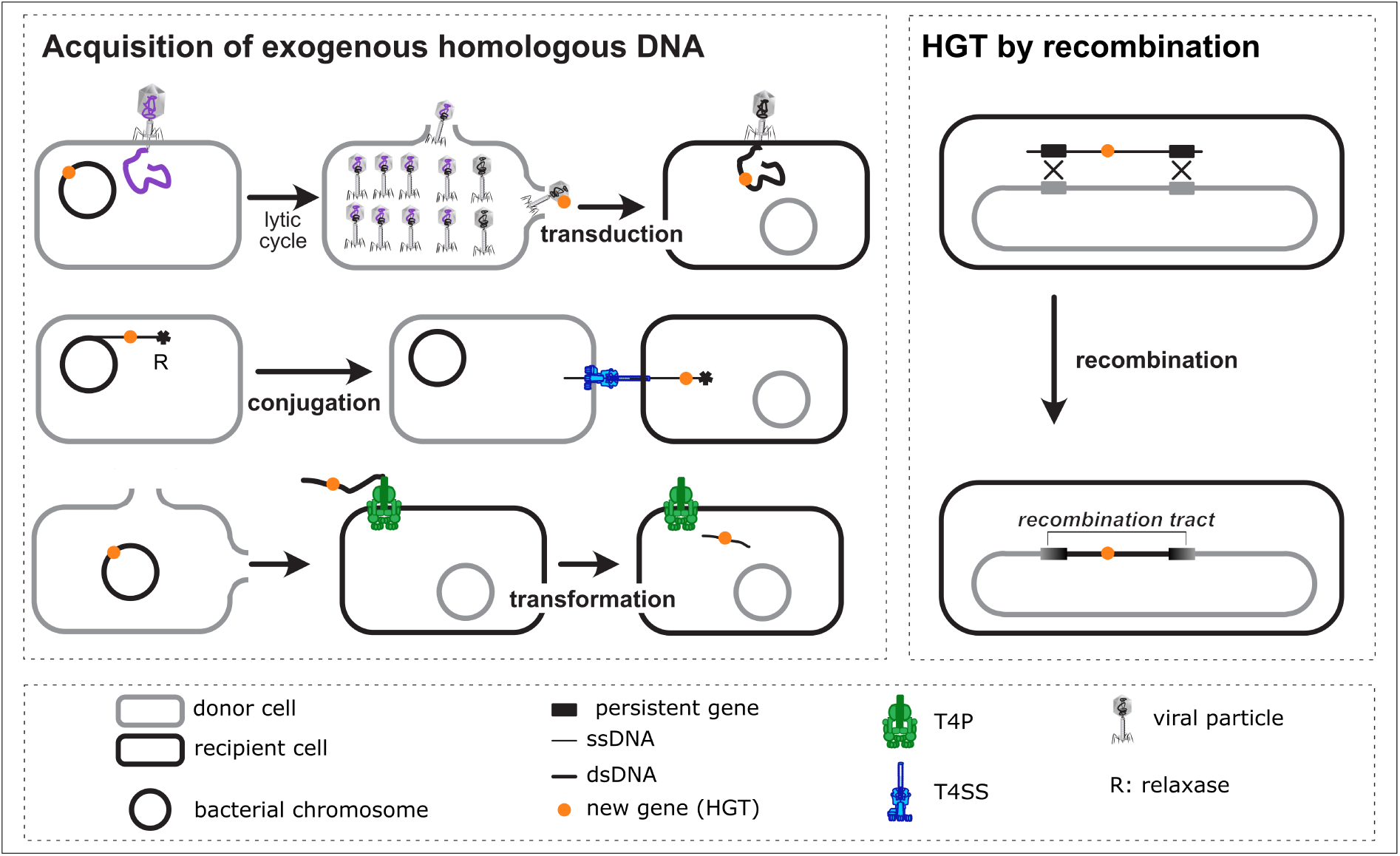
Different horizontal gene transfer mechanisms at the origin of homologous recombination tracts in a recipient genome. HGT: horizontal gene transfer; T4P: type IV pilus; T4SS: type IV secretion system; R: relaxase.

Natural transformation was proposed to provide several benefits. The uptaken DNA might be used as a source of nutrients for the recipient bacteria (11), as a substrate for repair of damaged DNA (12), or as a source of genetic variation by allelic recombination (13). The latter would favour the action of natural selection by reducing linkage disequilibrium (14), purging deleterious alleles (15), favouring the fixation of adaptive mutants (16), and curing chromosomes of costly mobile genetic elements (MGEs) (17). The latter process is favoured by the relatively small average size of exogenous DNA, which favours gene deletions over acquisitions of non-homologous DNA (17–19). This results in intragenomic conflict between the bacterium and the MGEs as the latter strive to defend themselves against deletion by blocking transformation (17). The average size of incoming DNA is crucial to the chromosome curing model (17). Transfer by natural transformation of genomic segments of different sizes (up to 150 kb) and carrying various functions has been observed in a wide variety of bacterial species in the laboratory (20–22). This process can result in the acquisition of small transposons (23,24), plasmids (25), integrons (24), and genomic islands (22,24,26,27). Even though the role of transformation in the acquisition of novel genes has often been described (24–27), its impact in bacterial populations is difficult to assess because the process only leaves one or two recombination tracts which are not easy to distinguish from the outcome of HGT by HFR-like conjugation (28), phage transduction (29), or other less-understood processes like outer-membrane vesicles (30) (Figure 1). Hence, the impact of natural transformation in gene acquisition in natural isolates has not been sufficiently detailed.

Even if the components required for natural transformation are encoded in many bacterial genomes, not all these bacteria could be shown to be transformable in the laboratory (31). Variations in the regulation of natural transformation could explain part of these negative results. Indeed, large variations in transformation rates have been observed between and within bacterial species (32–37). These variations could contribute to understand the impact of natural transformation on the acquisition of novel genes. For this, we previously obtained a large collection of genomes and transformation rates of two phylogenetically distant bacterial species *Legionella pneumophila* (Lp) and *Acinetobacter baumannii* (Ab) to study the genetic basis of such variations. We quantified transformation rates using a luminescence-based transformation assay corresponding to a gene gain (14). Our analyses showed that the sequence divergence between the recipient chromosome and the homology arms of their transforming DNA did not affect the patterns of transformation rates. It also showed independence of transformation rates and phylogenetic distances between strains, possibly because divergence was always low (<1%) (14). We observed large variations in transformation rates as well as frequent recent losses of transformability in both species and that transformation loss was counter-selected (14). This counterintuitive result could be explained by intragenomic conflicts between MGEs and natural transformation, in accordance with the view that the latter may cure the genome from the former (17). Hence, transformation could favour acquisition or loss of genes and the net overall effect remains unknown. Here, we try to disentangle the impact of natural transformation and other processes on the acquisition of novel genes by HGT.

Our hypothesis is that if natural transformation is important for the acquisition of novel genes, then the comparison between transformable and non-transformable strains should reveal it. For an accurate analysis, because transformation may result in the acquisition or loss of genes (20), this requires studying the patterns of recombination and transformation in light of the chromosome organisation and the types of genes that are acquired. We thus used the existing data on transformation rates in Ab and Lp to assess the impact of this process in the acquisition of novel genes. We evaluated the effect of transformation on gene gains and on the balance between gains and losses. We then estimated and characterised in several ways the effective contribution of natural transformation to the acquisition of novel genes by homologous recombination.

## Results

### Impact of natural transformation on gene flow and genome size

We analysed our previously published dataset of genomes and experimental determination of transformation phenotypes in 496 Ab strains and 786 Lp strains. Transformation was measured using a bioluminescence assay and varied over 6 orders of magnitude in Ab and 4 in Lp (14). Based on the experimental detection limits, we previously classed 64% of Ab and 52% of Lp strains as transformable and the rest as non-transformable (Figure 2, Table S1). Hence, we have a quantitative (TF^log^) and a qualitative (TF^bin^) measure of transformation. In parallel, we identified the pangenomes of both species, which contain 31,103 gene families in Ab and 11,932 in Lp. The persistent gene families were defined as those present in more than 95% of the genomes of each species. We identified 2629 (Ab) and 2325 (Lp) persistent genes that were used to compute recombination-free phylogenetic trees, as described before (14), which are used to infer ancestral states, make correction of statistical tests by phylogeny, and to identify terminal branches. Across this study we focus exclusively on terminal branches because the inference of ancestral states of transformation rates and identification of recombination tracts are less accurate for deeper regions of the tree (14).

**Figure 2.**
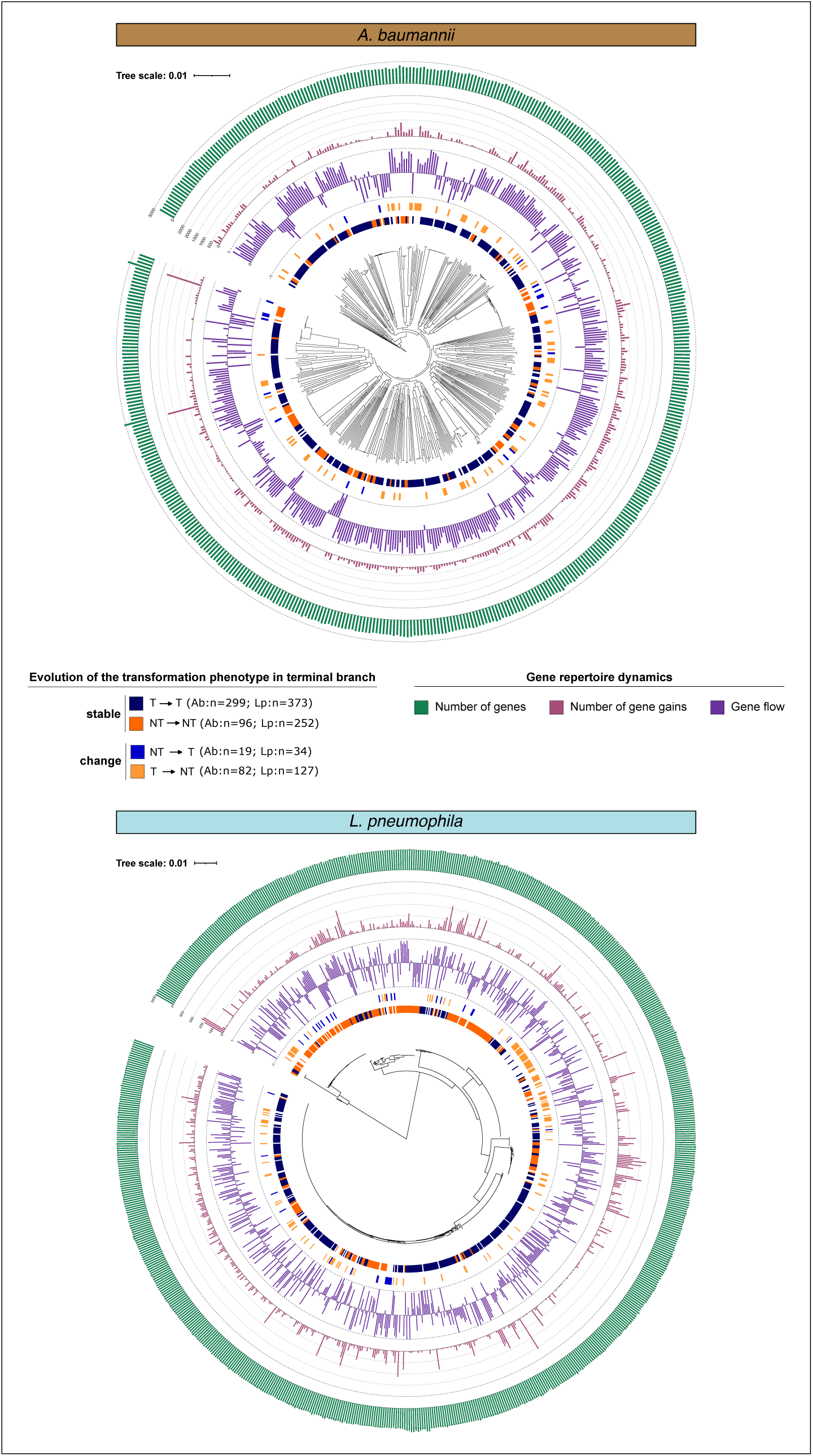
Recombination-free phylogenetic trees of the core genomes of *A. baumannii* (left) and *L. pneumophila* (right). The phylogenetic tree was retrieved from (*14*). The arcs around the trees represent (from inside to the outside). 1) the transformation phenotype of the strains whose transformation phenotype remained the same along the terminal branch (NT(ancestral)->NT(current), T(ancestral)->T(current)) 2) the transformation phenotype of the strains whose transformation phenotype changed along the terminal branch (NT(ancestral)->T(current), T(ancestral)->NT(current)) 3) net gene flow in terminal branch. 4) Number of gene gains in terminal branches. 5) Number of protein-coding genes in the genome.

We first tested the hypothesis that transformation is associated with differences in genome size, as measured by the total number of protein-coding genes. In Ab, the number of genes per genome was not significantly different between non-transformable and transformable strains when using the qualitative classification of transformation (TF^bin^) and was larger in transformable strains when using the quantitative phenotype (TF^log^) (Table S2: T1, T7). Yet, the latter effect was very low (R^2^ between 0.0061 and 0.0041) and at the edge of statistical significance (p=0.045). In Lp, transformable genomes were significantly smaller in all the variants of the analysis (p between 0.0015 and 10^-8^), but the trait also explained a very small fraction of the variance (R^2^ between 0.01 and 0.038).

We then used the pangenomes and the phylogenetic trees to infer past gains and losses of genes (Figure 2). We used this information to compute gene flow as the number of gains minus the number of losses, divided by their sum. The latter term allows to normalize by the total number of events, which are expected to depend on the branch length. Gene flow was significantly negative in all cases for both species (all statistical tests are in Table S2: see T4, T10), even though effects were low (R^2^ between 0.008 and 0.026 in Ab and between 0.045 and 0.028 in Lp). Finally, we analysed the specific association between gene gains and the transformation phenotype (Table S2). We found no significant association in Ab (Table S2: T2, T8, p>0.05) and a significant but weak negative association in Lp (Table S2: T2, T8, p<0.0001, R^2^≤0.03). In summary, transformation is weakly associated with a net loss of genes in genomes. In Lp, this is accompanied by a lower rate of gene gain whereas in Ab there is only a significantly higher rate of loss.

To gain a better understanding of the impact of transformability on gene flow we controlled more tightly for the effect of change in transformability (using the binary classification of transformability). We compared gene flow between strains predicted to have a non-transformable phenotype in the recent past (first parental node in the tree) and that have changed to become transformable (NT->T) or remained unchanged (NT->NT). In both species, the strains that became recently transformable had a more negative gene flow than those that remained non-transformable (Table S2: T24), the effect being small and only statistically significant for Ab (p<0.005). Altogether, these results suggest that transformation has a small negative impact on the net rate of acquisition of novel genes.

### Gene gains associated with transformability

During natural transformation the exogenous DNA integrates the chromosome by homologous recombination. To investigate gene gain potentially caused by natural transformation, we identified the tracts in the genomes that resulted from recent events of homologous recombination (in terminal branches). We previously showed that transformation is associated with elevated rates of recombination (14), and we now wished to test if genomic regions of high recombination rate coincide with regions with frequent gene gains as that might reveal gene gains by transformation. To compare regions across the genomes, which are highly variable in terms of gene repertoires, we used a method we have previously developed (3). We define *intervals* as the regions between two consecutive persistent genes in a genome. Hence, a gene gain can be placed in a specific interval of its genome. A set of intervals flanked by the same persistent gene families makes a *spot* (Figure 3A). Our past results showed that most spots are devoid of accessory genes because HGT is concentrated in a few ones (hotspots) (3). Accordingly, we identified 2612 spots in Ab among which only 588 had accessory genes. Around 80% of the latter had at least one recent gene gain. In Lp, we got 2315 spots among which only 449 had accessory genes and 50% had at least one gene gain. We then analysed the recent gene gains in the spots in relation to the presence of homologous recombination tracts in the flanking persistent genes (Figure 3B). This analysis reveals that there are tracts of homologous recombination across the chromosome, even if a few loci reveal higher rates. Intervals flanked by persistent genes covered by at least one recombination tract are positively associated with the presence of recent gene gains in both species and using any of the transformation phenotypes (Lp: χ^2^, Odd Ratio_gain/no gain_=14 in T and NT, p<2.2×10^-16^; Ab: χ^2^, Odd Ratio_gain/no gain_=5 in T and NT, p<2.2×10^-16^, see Methods). This positive association between recombination and recent gene gain suggest recombination in flanking persistent genes favours the acquisition of novel genes in both species.

**Figure 3.**
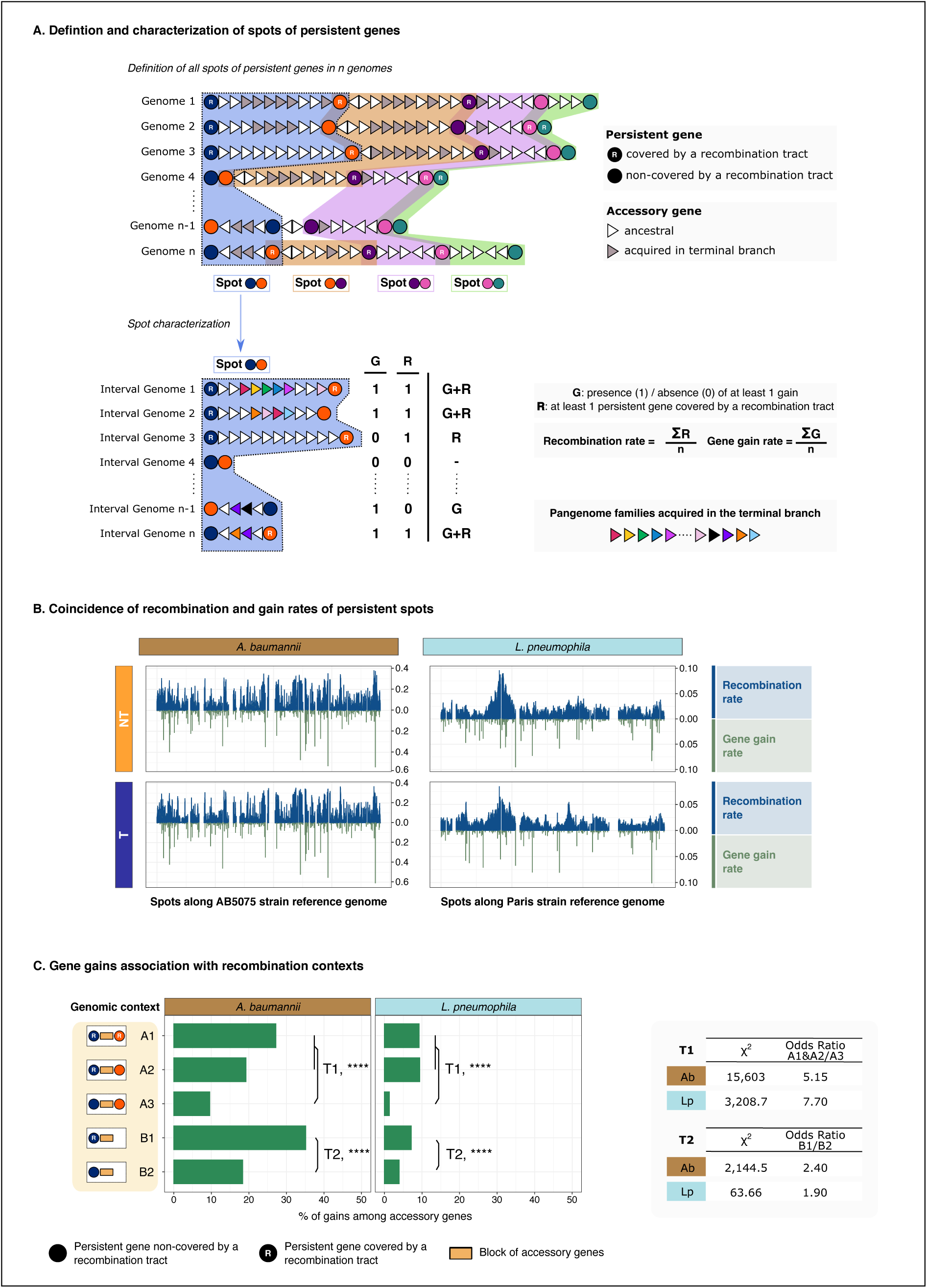
Analysis of gene gains in spots in relation to recombination in flanking accessory genes. **A. Scheme describing the definition of a spot of persistent genes in a collection of genomes and its characterization by the presence of gene gains and/or recombination tracts.** **B. Distribution of the recombination and gene gain rates at spots in relation to the transformation phenotype.** T: transformable, NT: non-transformable. **C. Distribution of the accessory genes in the different recombination contexts.** Statistical tests T1 and T2 are χ^2^ association tests. Odds ratios were calculated as the ratio of the fractions of the number of accessory genes in each genomic context that are gains over the ones that are not (see Methods). ****: p-value <2.2×10^-16^.

We then detailed the association between homologous recombination and gene gain while taking into account the existence of breaks in the genome assembly. For each accessory gene, we identified if it was gained in the terminal branch and assessed its recombination context (Figure 3 and S1): flanked by two (A1), one (A2; B1) or zero (A3; B2) recombining persistent genes. Genes in context A1, A2 and B1 were considered to potentially belong to the recombination tract of their neighbouring recombining persistent genes. While this may seem less obvious for A2 and B1, these account for cases where one of the edges of the recombination tract is in another contig (B1) or is not identifiable (A2). Also, events of illegitimate recombination in the non-homologous DNA (accessory genes) have been described and could result in patterns like A2 and B1 (27). Genes in contigs lacking persistent genes (C1 context) (Lp: 13% of the total; Ab:14% of the total) were excluded. The proportions of accessory genes identified as gains in A1 (Lp: 9.2%; Ab: 27%) and in A2 context (Lp: 9.4%; Ab: 19%) were not significantly different in Lp (χ^2^, p=0.87) and significantly lower in A2 compared to A1 in Ab (χ^2^, p<2.2×10^-16^), in agreement with the view that A2 may also account for recombination events. More importantly, both A1 and A2 revealed higher rates of gene gains than A3 in both species (Lp: χ^2^, p<2.2×10^-16^; Ab: χ^2^, p<2.2×10^-16^). When comparing the different contexts (A1&A2 vs A3 or B1 vs B2), we found more gene gains in the presence of recent recombination tracts (Figure 3C).

We then restricted our analysis to the gene gains associated with recombining persistent genes (A1, A2 and B1) and weighed them against all the gene gains that were inferred regardless of the recombination context. By doing so, and assuming that all these gains were caused by the identified recombination event, we got an upper limit estimate of the contribution of recombination to gene gains: 24.8% and 7.7% of all the gene gains in Ab and Lp respectively. Again, this suggests more acquisition of novel genes by recombination in Ab than in Lp.

As mentioned above, incoming DNA recombining with the chromosome may have been acquired by transformation or by other processes such as conjugation or transduction. To disentangle the contribution of transformation, we analysed the strains showing a stable transformation phenotype along the terminal branch leading to them, i.e., that had a binary classification of transformability similar to that of their most recent parental node in the species phylogenetic tree. In case of the strains that remained non-transformable, the observed recent gene gain caused by recombination, i.e., gains in A1, A2 and B1 contexts, should not be the result of natural transformation. It can thus be taken as an indication of the basal contribution of recombination by the other processes. The fraction of gene gains associated to recombination in these stably non-transformable strains is 17.3% in Ab and 5.9% in Lp. This can be compared with the rates of gene gains associated to recombination in stably transformable strains, which is 23.7% in Ab and 7.0% in Lp (Figure S2). If one assumes that the other sources of DNA are unchanged for strains capable of natural transformation, then one can estimate the share of natural transformation in the acquisition of novel genes in transformable strains by making the difference with the average values for the strains that remained non-transformable (see Discussion, Figure S2). This would suggest that gene gains due to natural transformation are 6.4% of the total in Ab and 1.1% in Lp, which would represent 26.8% (Ab) and 16.4% (Lp) of the novel genes acquired by recombination in these species.

### Transformation favours fewer and shorter gene acquisition events

Gene gain by transformation is expected to be associated with events implicating relatively few genes, when compared with the impact of phages and conjugative elements. This is because entry of the transforming DNA in the cell is accompanied with the action of endonucleases that create relatively small fragments (38,39). In contrast, viral particles transduce DNA fragments of the size of the phage genome (often around 50 kb for temperate phages) and conjugation can transfer entire chromosomes. To understand if transformation might be associated with acquisition of small numbers of genes, we identified the individual events of gene gain and loss. We clustered into one *event* the arrays of genes gained in the same terminal branch and present between the two same persistent genes (A context) or between the same persistent gene and the end of a contig (B context), when they have few or no other gene in between them (see Methods, Figure 4A). The allowance of some gene families classed as not recently acquired is to account for transposition of insertion sequences or gene families at high copy number. Yet, these interruptions remained exceptional since most events have none in Ab (85%) and in Lp (91%) (Figure S3). We counted the number of acquisition events per genome and compared it between the two phenotypes (T and NT) while taking into account the phylogeny. In Ab, the number of acquisition events was significantly and positively associated with transformable strains when using both the quantitative (Table S2: T11, p=0.037) and the qualitative phenotype (Table S2: T5, p=0.035) even though the transformation trait explained little of the variance (R^2^=0.028). In Lp, no significant association was found between the transformation phenotype of a strain and the number of acquisition events we observed in its genome (Table S2: T5, T11).

**Figure 4.**
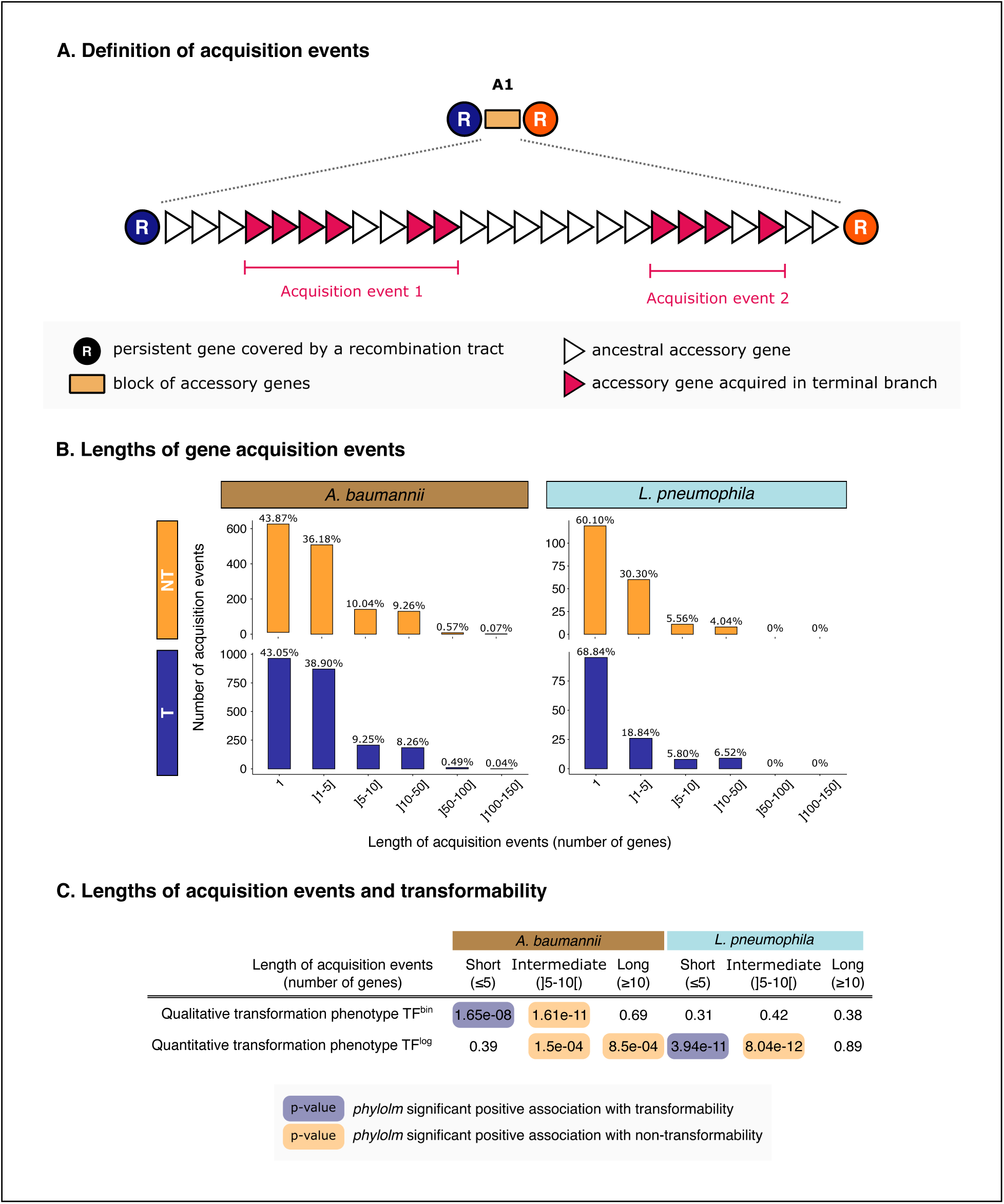
Characterization of the gene acquisition events by recombination. **A. Definition of gene acquisition events between two recombining persistent genes.** **B. Distribution of the lengths of gene acquisition events per genome depending on the transformation phenotype in *A. baumannii* and *L. pneumophila*.** **C. Average proportion of gene acquisition events of different lengths depending on the transformation phenotype in *A. baumannii* and *L. pneumophila* and their association with transformability.**

The acquisition events had between 1 and 107 genes in Ab and between 1 and 41 genes in Lp (Figure 4B). The average length of acquisition events was longer in Ab compared to Lp (Ab: 4.4 genes; Lp: 2.8 genes). To detail these observations and explore the possible relation between event length and transformation phenotype, we defined three categories of acquisition events: short (≤5 genes), intermediate (5 to 10 genes) and long (≥10 genes) and compared their proportions among the events observed in each genome (Figure 4C). Short events were positively associated with transformability in Ab and Lp (Table S2: T25, T27), even if for Lp this was only significant for the quantitative transformation phenotype and for Ab for the qualitative phenotype (Figure 4C). Expectedly, intermediate-size and long gene acquisition events followed the inverse trends. In Ab, both were associated with non-transformability whether we used the qualitative or the quantitative transformation phenotype (Figure 4C, Table S2: T26, T27, T29, T30). In Lp the intermediate-size events were also significant with the quantitative phenotype (Figure 4C, Table S2: T29), but we found no statistical signal for long events possibly because these are quite rare and the statistical test may lack power. Hence, transformable strains tend to have an excess of events with less than 6 genes and a lack of those with more than five genes. These results are thus in agreement with the hypothesis that transformation favours events of acquisition of a small number of genes.

### Exchanges in hotspots of gains

Spots with many gene gains (hotspots) can result from the integration of MGEs by site-specific recombination or gene acquisition by homologous recombination. Since only the latter concern transformation, hotspots could differ between transformable and non-transformable strains. We separated transformable and non-transformable strains and defined the sets of unique gene families that were gained in each spot. We defined hotspots as the minimal set of spots necessary to accumulate 90% of the gene families gained across all spots (see Methods). The other spots were deemed coldspots. Hotspots represented only 3.48% of all the spots in Ab and 6.05% in Lp. They are scattered along the chromosomes of both species (Figure S4). We then identified the hotspots common to the sets of transformable and non-transformable strains in each species. Two thirds of the hotspots were similar in the two sets of Ab while only around 30% were similar in Lp (Figure S4). The specificity of some hotspots for one phenotype could be explained by a small number of genomes having this spot and harbouring gene gains thus decreasing the likelihood of having it represented in both phenotypes. This is exactly what we observed: in both species, the hotspots specific to transformable and non-transformable strains had on average 3 times fewer genomes with gene gains than the hotspots common to both phenotypes. All the next results are qualitatively similar when performing the analyses with the hotspots defined on each group or on all genomes regardless of the transformation phenotype (Figure 5 and S5). In both species and for both transformation phenotypes, hotspots of gains had a higher recombination rate at the flanking persistent genes than coldspots (Lp:Wilcoxon, p=4.5×10^-5^; Ab: Wilcoxon, p=3.9×10^-7^) (Figure 5A). Within hotspots, we collected the intervals with gene gains and then separated them in two groups: those with and those without evidence of homologous recombination. The majority of intervals in hotspots with recent gains lacked flanking recombining genes (Figure 5B) in both transformable and non-transformable strains. Hence, large integration events are more frequently flanked with recombining persistent genes, but still most integration events in these loci are not associated with homologous recombination.

**Figure 5.**
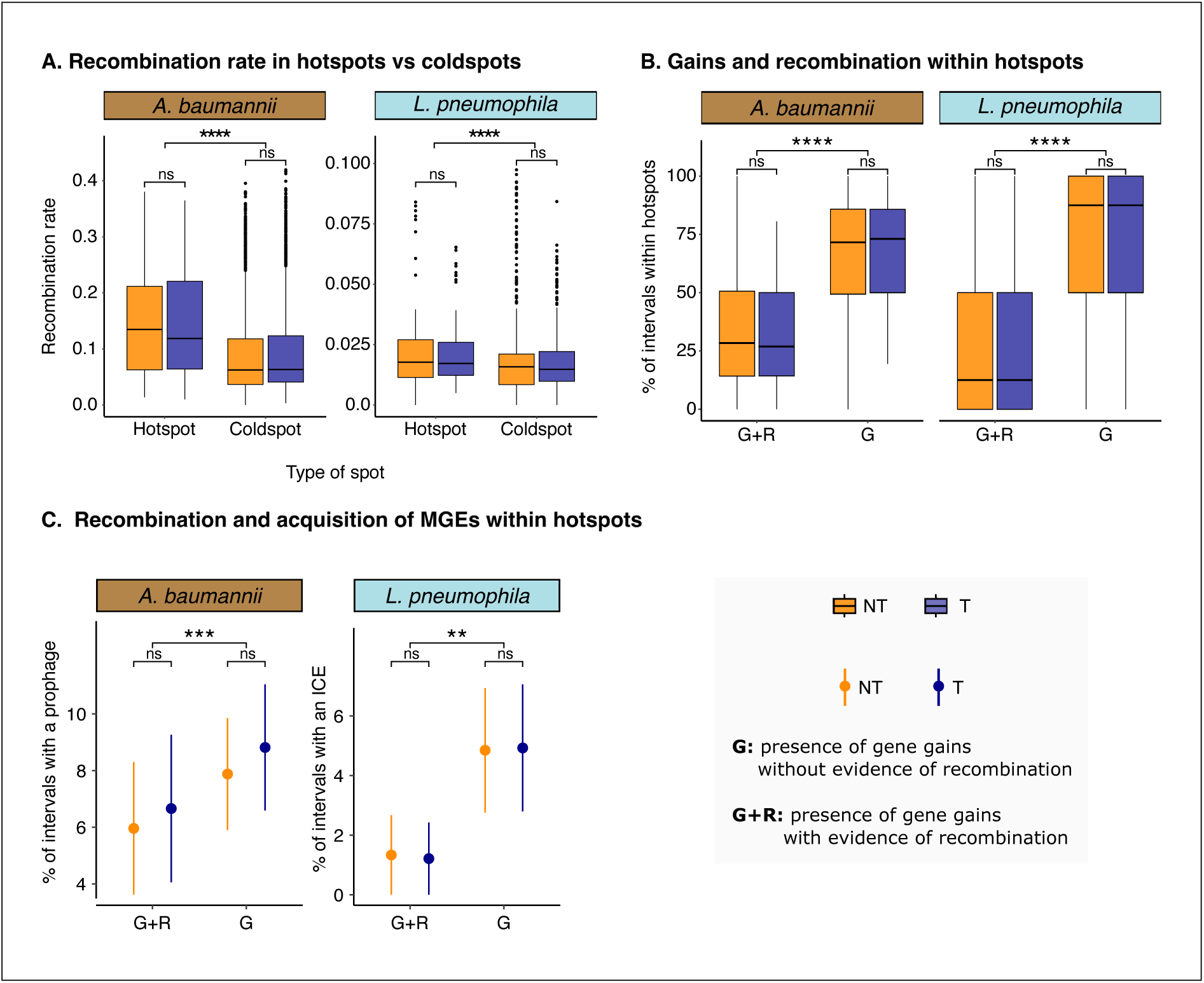
Characterization of hotspots of gene gains. **A. Distribution of recombination rates in persistent genes flanking hotspots and coldspots depending on the transformation phenotype.** **B. Comparison of the proportion of intervals with at least one gene gain in hotspots.** The data is stratified in terms of the evidence for recombination in neighboring persistent genes in terms of the transformation phenotype. **C. Proportion of intervals with recently acquired prophage in Ab and with recently acquired ICE in Lp within hotspots.** The data is stratified in terms of the evidence for recombination in neighboring persistent genes of the intervals considered and in terms of the transformation phenotype. Dots stands for the average proportions and bars for the standard error. Statistical tests are Wilcoxon tests with ***: p<0.001, **: p<0.01.

These results are consistent with the idea that most large events of gene acquisition are of MGEs encoding their own integrases.To confirm this hypothesis, we characterized the large MGEs in both species, especially integrative conjugative elements (ICEs) and prophages. Conjugative systems were more frequent in Lp (811 conjugative systems in 699 genomes out of 786) than in Ab (13 conjugative systems in 13 genomes out of 496) (Figure S6). Prophages were absent from Lp, as previously described (14), and were frequent in Ab (1215 elements), where 88% of the strains carried at least one element (Figure S7). As expected, intervals with gene gains which had no evidence of homologous recombination in flanking persistent genes were more often the sites of integration of large MGEs than the ones with homologous recombination at the flanking persistent genes (Figure 5C). This was true for hotspots in transformable strains (Lp:Wilcoxon, p=0.015; Ab:Wilcoxon, p=0.0062) and in non-transformable strains (Lp:Wilcoxon, p=0.014; Ab:Wilcoxon, p=0.0038). This confirms the hypothesis that integration of the large MGEs tends to be regrouped in a few loci in the chromosome, the hotspots, by site-specific recombination rather than homologous recombination and are thus not usually acquired by natural transformation.

HGT drives the acquisition of antimicrobial resistance (AMR) genes, raising the hypothesis that natural transformation is involved in this process (40,41). We identified AMR genes in both species and estimated their proportion brought by recombination. We focused on the AMR genes which were not persistent and thus susceptible to have been brought by any HGT process. In Ab, 1515 accessory genes were marked as AMR. These correspond to 69 gene families of the pangenome and 13 different AMR classes. In Lp, we identified 1781 genes corresponding to only 3 gene families and 2 AMR classes. We then searched for these genes among the events of gene acquisition by homologous recombination identified above, that is to say in context A1, A2 and B1. In Lp, no AMR gene was recently acquired in the terminal branch and the analysis couldn’t be done. In Ab, 9.6% of the AMR genes were gained in the terminal branch. 14% of them, mainly fosfomycin and aminoglycoside resistance genes, were found within the acquisition events we identified above and thus acquired by homologous recombination. None of the acquisition events carrying these AMR genes were found in phagesor integrative conjugative elements and around 86% of them were shorter than 5 genes. Some could hence result from transformation events. Many AMR in *A. baumannii* are known to accumulate in the AbaR island (42). Unfortunately, this locus cannot be analysed with the current dataset, because the large number of transposable elements breaks contig assemblies complicating the analysis of the spots (context C in Figure S1).

## Discussion

Throughout this study we aimed at quantifying the effect of natural transformation on the acquisition of novel genes. Evaluating this effect requires to identify the imprint of recombination in bacterial genomes, infer the recombination events resulting in gene gains, and then assign these gains to transformation. All these tasks are challenging. Recombination is intrinsically difficult to measure because recombination tracts are difficult to pinpoint when exchanges take place between very similar sequences and because multiple overlapping events of recombination produce complex tracts. We used the information on recombination at flanking persistent genes to assess the likelihood that in between these persistent genes accessory gene gains were caused by recombination. We assumed that all recent gene gains flanked by recombination tracts were caused by homologous recombination. This should be seen as an over-estimate since recombination and gene gain may occasionally have occurred independently. This over-estimate is expected to be larger when only one of the flanking persistent genes is identified as recombining, and indeed cases A2 show a slightly smaller effect than A1 in figure 3C, even if both are significantly higher than cases A3. As datasets with completely assembled genomes become bigger, one will be able to better details the chronologies of gene gains and recombination to confirm how frequently the latter results in the former. As mentioned above, gene gains by recombination may arise from other processes such as conjugation and transduction. Unfortunately, there is no specific hallmark allowing to distinguish between these different sources of homologous recombination and the concentration of accessory genes in hotspots of MGEs makes disentangling the different contributions particularly hard. Still, by comparing transformable and non-transformable bacteria, we were able to leverage the differences in recombination and gene gain to propose a first estimate of the impact of transformation in the acquisition of novel genes. This estimate is relatively low (6% in Ab and 1% in Lp) suggesting that transformation is responsible for the acquisition of a minor fraction of the genes in the genome.

The effect of natural transformation on the bacterial gene repertoires might be reflected in the genome size of bacteria. Despite the potential changes in gene repertoires caused by natural transformation, the gene repertoire sizes in Ab and Lp were similar between transformable and non-transformable strains. Contrary to *Aggregatibater actinomycetemcomitans* (43) and *Streptococcus pneumoniae* (17) whose non-competent strains had smaller genomes than the competent ones, genome size was not much affected by bacterial transformability in our much larger datasets on Ab and Lp. The lack of strong effect of transformation on genome size could be explained by novel genes brought by natural transformation being compensated by additional MGE losses by chromosome curing. Transformable bacteria acquire some genes by transformation but might effectively acquire fewer genes by MGE (because the latter are quickly deleted by transformation). In contrast, non-transformable bacteria will not get novel genes by transformation but the increased stability of the acquired MGEs compensates for this absence. In both species, gene gains and gene losses were not exactly balanced and transformable bacteria showed a net loss of genes in genomes. In strains that recently became transformable, this net loss of genes could be related to the loss of transformation-inhibiting MGEs. This complex interplay between two sources of gene gain, transformation and MGE-mediated, might lead to the small net negative effect of transformation on gene flow. But this net loss of genes was small enough not to significantly affect the genome size.

Gene acquisition events by homologous recombination differ between transformable and non-transformable strains in number and in length. In Ab, transformable strains had more acquisition events than the non-transformable ones. The difference was not significant for Lp. In both species, however, transformable strains were associated with short acquisition events while their non-transformable counterparts were associated with longer ones. Previous experimental studies in *S. pneumoniae* (44)*, Helicobacter pylori* (45) and *Haemophilus influenzae* (35) showed that most recombination events observed during transformation are short. In a parallel evolution experiment in *Bacillus subtilis*, transferred segments between two *B. subtilis* lineages contained an average of 5.1 genes and had an exponential length distribution (46). The overrepresentation of short length fragments in transformation events may be due to the action of endonucleases during the DNA entry in the cell (38). After its uptake during replication, the transforming DNA, especially heterologous region flanked by homologous sequences, can also be exposed to digestion by restriction endonucleases from restriction-modification systems (47). This overrepresentation of short length fragments could be reinforced by the quality of DNA available to the transformable bacteria, since exogenous DNA may often be fragmented in the environment thus providing short DNA fragments rather than very large ones. The experiment in *B. subtilis* also showed that the transferred segments were distributed over the entire core genome of the recipient (46). In agreement with this observation, we found that long events of gene acquisition are concentrated in few regions and related to the presence of large MGEs such as phages in Ab and ICEs in Lp whereas gene gains by homologous recombination tend to be more scattered in the chromosome. Hence, transformation may result in more homogeneous recombination rates across the chromosome.

Despite the difficulty in disentangling the effect of transformation from all the other HGT processes that may act in recombination regions, we propose a first estimate of the contribution of natural transformation to the acquisition of novel genes. Our method relies on the assumption that the basal contribution of other mechanisms to recombination, conjugation and transduction, is the same in transformable and non-transformable strains. One might think that this results in an under-estimate of the effect of transformation because transformable strains have fewer MGEs. Yet, the MGEs driving conjugation and transduction are not in the recipient cell, they are in the donor, which may or may not be transformable and may or may not have many MGEs. Hence, the effect of chromosome curing of MGEs in recipient cells may have a very minor effect in our estimate. Both species are known to recombine at significant rates (48–50), but the impact of recombination in gene gain seems higher in Ab. Future work will be needed to assess how this is affected by sampling strategies, which were very different for both species (14). Interestingly, among genes acquired by recombination, the estimated contribution of transformation is similar between the species. Our estimates suggest that transformation does play a significant role in the acquisition of novel genes albeit smaller compared to with MGE-mediated HGT processes.

The percentage of genes acquired by transformation does not suffice to evaluate the evolutionary importance of this process. Transformation does not involve MGEs and may result in the acquisition of different types of functions with diverse adaptive potential. The acquisition of a novel capsule operon by transformation in Griffith’s experience leading to a change in virulence is a remarkable example of that (51). Similar serogroup conversions by transformation were observed in *Vibrio cholerae* (52). Other functions, such as enzymes involved in metabolism were found in *V. cholerae* (53) and *Campylobacter coli* (54). Our results suggest that almost fifteen percent of the recently acquired AMR genes arrived by recombination independently from MGEs. This supports the idea that natural transformation plays a role in the dissemination of antimicrobial resistance (40). Transfer of clinically relevant AMR genes by natural transformation was observed in *Enterococcus faecalis* (55), *Haemophilus influenza*, *Haemophilus parainfluenzae* (56), *Streptococcus pneumoniae*, *Streptococcus oralis*, *Streptococcus mitis* (57), *Neisseria meningitidis* (58), and *A. baumannii* (22,59,60). These transfers may occur between species of the same genus, e.g. the large antibiotic resistance islands AbaR can be transferred by transformation across pathogenic *Acinetobacter* species with down to 90% sequence identity (22). In spite of the experimental data showing that transformation can spread AMR in the laboratory, evidence in clinical isolates has been missing. This is partly due to the difficulty in assigning the source of recombination-driven gene transfer to a specific HGT mechanism, including transformation. Further work is required to systematically disentangle the impact of the different mechanisms of transfer on the observed patterns of gene gain by recombination.

## Material and Methods

### Collection of Acinetobacter baumannii and Legionella pneumophila strains

We studied a collection of 496 environmental and clinical strains of Ab and another collection of 786 clinical strains of Lp (14). Here, we provide a short summary of these methods, which can be found in our previous publication (14). We analysed the genome assemblies to define the sequence types (STs), pangenomes, phylogenies and recombination tracts of these strains in both collections. Draft assemblies were annotated with *prokka* (61) called from *PanACoTA v.1.2.0* (62). Both pangenomes were built using single-likage clustering so as to form pangenome families with *PanACoTA*. Pangenome families are sets of proteins sharing at least 80% identity. We defined a pangenome family as a persistent gene family when at least 95% of the genomes had a unique member of the family. Recombination-free phylogenies were built on the concatenated alignment of persistent genes ordered according to a reference genome with *IQTree v.1.6.12 modelfinder* (best-fit model: TVM+F+I+G4; 1,000 ultrafast bootstraps) and were rooted with an outgroup species. The recombination tracts were identified from the alignments of persistent genes using *Gubbins v.2.4.1* (63). While Gubbins is originally intended to analyse only closely related genomes, a recent study showed it was equivalent to other state-of-the-art approaches in identifying recombination, that don’t scale to datasets of this size, when one focus on recent events (as is the case in this work) (64). A discussoin on these issues can found in our previous publication (14). Transformation rates of the strains were measured with a transformation luminescence assay and the strains were categorized as transformable or non-transformable based on the transformation rate of known non-transformable strains (Table S1). The transformation phenotype across the phylogenetic tree were previously reconstructed using *PastML v.1.9.34* (65) but this reconstruction was, here, only of interest in the terminal branches. We also thoroughly characterized the diverse mobile genetic elements present in the genomes notably integrative conjugative elements (ICEs) reduced to conjugative systems present on contigs that were not of plasmid origin (*Macsyfinder 20221213.dev* (66,67)), prophages (*VirSorter v*.*2*.*2*.*3* (68)*, CheckV v*.*0*.*7*.*0* (69)) and insertion sequences (*ISEScan v*.*1*.*7*.*2*.*3* (70)). As described in (14), the contigs that were of plasmid origin were identified using their weighted gene repertoire relatedness (wGRR) with plasmids allowing us to distinguish conjugative systems of ICEs from the ones of plasmids.

### Identification of antimicrobial resistance genes

Antimicrobial resistance genes were searched in the genomes of Lp and Ab with *AMRFinderPlus v.3.12.8* (71) (default parameters).

### Ancestral reconstruction of pangenome families presence/absence

We represented the pangenome as a binary matrix of presence or absence of each gene family in each genome. We reconstructed the ancestral states of this pangenome in terms of presence-absence in each node of the recombination-free rooted phylogenetic tree. This was done using the MPPA prediction method and the F81 evolutionary model of *PastML v.1.9.34* (65). Since the MPPA algorithm can keep several ancestral states per node if they have similar and high probabilities, we only kept the events where both ancestral and descendant nodes had one single distinct state either presence or absence. Using this method, on average per node more than 99% (Lp:99.8%; Ab:99.7%) of all the pangenome families had a confident presence/absence state in both species.

### Measure of gene gains, gene losses and gene flow in terminal branches

We focused on recent gains and losses of pangenome gene families, i.e., on events that were inferred to have occurred in the terminal branches of the species trees. A *gene gain* is defined as a gene family that is present in the leaf but inferred to be missing in the parental node. A *gene loss* is defined as a gene family that was inferred to be present in the parental node but missing in the leaf. We defined *gene flow* of a strain as the balance between gene gains and losses:

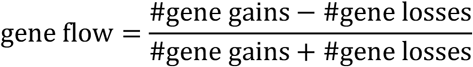

This ratio provides a symmetric measure of gene flow bounded between -1 and 1. The division by the sum of gains and losses allows to normalize the gene flow across branches that have different sizes and thus naturally different numbers of events. If the gene flow is negative then gene losses are more prominent than gains. If it is positive, then bacteria favour gene gains over losses. Close to 0, gene gains and losses are approximately in similar numbers.

### Measure of the recombination and gene gain rates of a spot

We use persistent genes to identify the location of gene acquisitions. Genomes are split in a collection of intervals, which are defined as a succession of pairs of persistent genes. Hence, each interval in each genome is a region comprised between two consecutive persistent genes of a same contig. Since persistent genes are present in most genomes, one can group intervals from different genomes based on these persistent genes. We define a spot as a collection of intervals (typically one per genome), that are flanked by the same two families of persistent genes and present in at least 50% of the genomes of the collection. We consider the interval as recombining in a strain when at least one of the two flanking persistent genes is part of a recombination tract. This simplification comes with limitations. In the case of the two flanking persistent genes being part of the recombination tract, it is likely that the genes comprised in the interval were also part of the recombination tract. However, when only one of the two flanking persistent gene belongs to a recombination tract, it is harder to precisely delimit the recombination tract. This could lead to overestimation of the number of gains obtained by recombination. An interval harbouring one or several gene gains in a strain was considered having at least one acquisition event (see Identification of gene acquisition events and Figure 3). For every interval in each strain, we consider two binary variables: presence/absence of an acquisition event (G) and presence/absence of a recombination event (R) in the flanking genes. We then summarized these information on all strains which had the spot. We were thus able to calculate a recombination rate and a gene gain rate for every spot (Figure 2A). We defined them as the number of times an interval of the spot was observed as having a recombination or acquisition event divided by the number of genomes present in this spot, respectively. The odds ratio of an interval with recombination evidence having an acquisition event were calculated as follows:

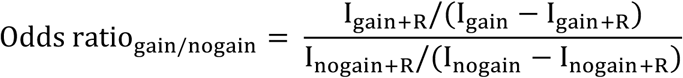

With I_gain_ the total number of intervals with at least one acquisition event, I_gain+R_ the number of intervals with at least one acquisition event and with evidence of recombination, I_nogain_ the total number of intervals without any acquisition event and I_nogain+R_ the number of intervals without any acquisition event but with evidence of recombination.

Of note: since our analysis of recombination and gene acquisitions is made only on terminal branches, each event is only counted once (no phylogenetic dependency).

### Definition of the recombination context of any accessory gene

An accessory gene can be in three different types of genomic contexts regarding persistent genes (Figure S1): in an interval delimited by two persistent genes (A), in a genomic segment delimited by only one persistent gene and the end of a contig (B) or in a contig lacking persistent genes and thus outside of any interval (C). Depending on the recombination state of the neighbouring persistent genes, we can distinguish several subcontexts. In context A, the two flanking persistent genes are part of a recombination tract (A1), only one of the two flanking persistent genes is (A2) or none of them is (A3). We can do the same for context B: the neighbouring persistent gene belongs to a recombination tract (B1) or not (B2). All the counts of accessory genes in each recombination context are to be found in Table S3. The odds ratios of an accessory gene being a gain in certain genomic contexts shown in Figure 2 were calculated as follows:

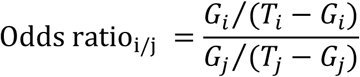

With G_i_ and G_j_ being the number of gene gains in genomic contexts i and j repectively and T_i_ and T_j_ the total number of accessory genes in genomic contexts i and j respectively.

### Estimation of transformation contribution to gains

Among all the accessory genes that were gained (G), the ones in context A1, A2 and B1 were considered to be acquired by recombination (G^REC^). Indeed, the recombination tract covering the persistent gene in context A2 and B1 could extend further than the bordering persistent gene due to events of illegitimate recombination in the heterologous part of DNA and thus encompass these accessory genes. We focused on stable non-transformable strains (NT>NT), i.e., strains that remained non-transformable from the parent node to the leaf in the terminal branch. In these strains, gene gains should not be the result of a transformation event. Hence, the proportion of gene gains acquired by recombination in stable NT strains corresponds to the basal contribution of recombination to the acquisition of novel genes in the absence of transformation (presumably occurring by transduction or conjugation). One can then estimate the contribution of transformation to gene gain (G^transformation^) by subtracting this basal contribution of recombination to the proportion of recombination-acquired gains in the transformable strains that remained transformable from the parent node to the leaf in the terminal branch (Figure S2):

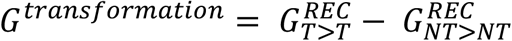

### Identification of gene acquisition events

The set of genes gained in an interval, i.e., between two consecutive persistent genes (context A1 and A2) or between one persistent gene and the end of a contig (context B1), may have been acquired in one or multiple events. To cluster gene gains in acquisition events, we had to choose a maximal number of genes separating two successive gene gains that belong to a single acquisition event. We made the choice of 3 genes based on several arguments. 1) By separating acquisition events by a number of accessory genes less than the average gap length (3.5 genes, Figure S8) between two successive gene gains, we avoided underestimating the number of acquisition events by clustering too many gene gains together while they were coming from different acquisition events. 2) The maximum number of genes separating two gene gains from the same acquisition event accounts for the disruption caused by insertion sequences (IS). Most IS (Lp: 99.8%, Ab:95.6%) that interrupts acquisition events are of a length less or equal to 3 genes (Figure S9). Transposable elements often have identical homologs in the genome and in that case they are not marked as gains. Our procedure allows to avoid multiplying unduly the number of events because of these elements. In every interval with gene gains in the terminal branch, we then clustered successive gene gains separated by less than 3 accessory genes into single gene acquisition events (Figure 3). In most intervals only one acquisition event was observed (Figure S10). The length of each acquisition event was measured as the number of genes (gains and genes in between) that are part of it. We classified the acquisition events in three different categories: *short* for when including less than 5 genes, *intermediate* for those having between 5 and 10 genes and *long* for those with more than 10 genes.

### Estimation of the coincidence between recombination and hotspots of gene gains

For each spot we listed the set of unique pangenome families that was gained at least once across the genomes considered. We then ranked the spots from the biggest set size to the smallest. The bigger the set of a spot is, the higher the diversity of the acquired gene families is. We browsed the spots in the previously defined order and at each spot, we counted the number of unique pangenome families that had been gained in this spot but not in the spot before. We then cumulated these numbers according to the previous spot ranking (Figure S11). Spots with the biggest set size and whose cumulated set of unique gains constitute 90% of the total number of unique pangenome families gained across all spots were considered as hotspots of gene gains (Figure S11). The gene content of these hotspots is more dynamic and encline to mobility as shown by the high diversity in gene gains they harbour. All the spots that were not categorized as hotspots were classified as coldspots. In each species, we applied this method by first considering all genomes and then each subcollection of transformable and of non-transformable genomes. These hotspots where then analysed in relation to recombination. We verified that working on all genomes or on each subcollection gave consistent results. We only present the results of each subcollection in the main results. The results obtained on all genomes can be found in the supplementary materials (Figure S5).

### Strain specificities affected by transformation rates

To take into account the phylogenetic structure of the bacterial population in our statistical analyses of the impact of the transformation trait on other traits across the tree, we used phylogenetic linear models (*phylolm* function form *phylolm* R package, default parameters) (72). We expressed different traits such as the number of genes, the gene flow or the number of acquisition events as a function of the transformation phenotype. The latter was used either as a qualitative (TF^bin^) or a quantitative (TF^log^) phenotype. All the statistical tests were listed in Table S2.

### Graphical representations

All graphs were produced with *R v.4.1.0*.

## Acknowledgements

This work was supported by the INCEPTION project [PIA/ANR-16-CONV-0005], the Laboratoire d’Excellence IBEID Integrative Biology of Emerging Infectious Diseases [ANR-10-LABX-62-IBEID] and the grant TransfoConflict [ANR-20-CE12-0004]. This work used the computational and storage services (TARS cluster) provided by the IT department at Institut Pasteur, Paris.

## References

1. Huang M, Liu M, Huang L, Wang M, Jia R, Zhu D, et al. The activation and limitation of the bacterial natural transformation system: The function in genome evolution and stability. Microbiol Res. 2021 Nov 1;252:126856.

2. Johnston C, Martin B, Fichant G, Polard P, Claverys JP. Bacterial transformation: distribution, shared mechanisms and divergent control. Nat Rev Microbiol. 2014 Mar;12(3):181–96.

3. Oliveira PH, Touchon M, Cury J, Rocha EPC. The chromosomal organization of horizontal gene transfer in bacteria. Nat Commun. 2017 Oct 10;8(1):841.

4. Torrance EL, Diop A, Bobay LM. Homologous recombination shapes the architecture and evolution of bacterial genomes. Nucleic Acids Res. 2024 Dec 24;gkae1265.

5. Griffith Fred. The Significance of Pneumococcal Types. J Hyg (Lond). 1928 Jan;27(2):113– 59.

6. Avery OT, MacLeod CM, McCarty M. STUDIES ON THE CHEMICAL NATURE OF THE SUBSTANCE INDUCING TRANSFORMATION OF PNEUMOCOCCAL TYPES. J Exp Med. 1944 Feb 1;79(2):137–58.

7. Blokesch M. Natural competence for transformation. Curr Biol. 2016 Nov 7;26(21):R1126– 30.

8. Zuke JD, Burton BM. From isotopically labeled DNA to fluorescently labeled dynamic pili: building a mechanistic model of DNA transport to the cytoplasmic membrane. Microbiol Mol Biol Rev. 2024 Mar 11;88(1):e00125–23.

9. Dubnau D, Blokesch M. Mechanisms of DNA Uptake by Naturally Competent Bacteria. Annu Rev Genet. 2019;53(1):217–37.

10. Lorenz MG, Wackernagel W. Bacterial gene transfer by natural genetic transformation in the environment. Microbiol Rev. 1994 Sep;58(3):563–602.

11. Redfield RJ. Genes for breakfast: the have-your-cake-and-eat-it-too of bacterial transformation. J Hered. 1993;84(5):400–4.

12. Redfield RJ. Evolution of Natural Transformation: Testing the DNA Repair Hypothesis in Bacillus Subtilis and Haemophilus Influenzae. Genetics. 1993 Apr;133(4):755–61.

13. Ambur OH, Engelstädter J, Johnsen PJ, Miller EL, Rozen DE. Steady at the wheel: conservative sex and the benefits of bacterial transformation. Philos Trans R Soc B Biol Sci. 2016 Oct 19;371(1706):20150528.

14. Mazzamurro F, Chirakadavil JB, Durieux I, Poiré L, Plantade J, Ginevra C, et al. Intragenomic conflicts with plasmids and chromosomal mobile genetic elements drive the evolution of natural transformation within species. PLOS Biol. 2024 Oct 14;22(10):e3002814.

15. Treangen TJ, Ambur OH, Tonjum T, Rocha EP. The impact of the neisserial DNA uptake sequences on genome evolution and stability. Genome Biol. 2008 Mar 26;9(3):R60.

16. Cavassim MIA, Andersen SU, Bataillon T, Schierup MH. Recombination Facilitates Adaptive Evolution in Rhizobial Soil Bacteria. Mol Biol Evol. 2021 Dec 1;38(12):5480–90.

17. Croucher NJ, Mostowy R, Wymant C, Turner P, Bentley SD, Fraser C. Horizontal DNA Transfer Mechanisms of Bacteria as Weapons of Intragenomic Conflict. PLOS Biol. 2016 Mar 2;14(3):e1002394.

18. Apagyi KJ, Fraser C, Croucher NJ. Transformation Asymmetry and the Evolution of the Bacterial Accessory Genome. Mol Biol Evol. 2018 Mar;35(3):575–81.

19. Tuffet R, Carvalho G, Godeux AS, Mazzamurro F, Rocha EPC, Laaberki MH, et al. Manipulation of natural transformation by AbaR-type islands promotes fixation of antibiotic resistance in Acinetobacter baumannii. Proc Natl Acad Sci. 2024 Sep 24;121(39):e2409843121.

20. Blokesch M. In and out—contribution of natural transformation to the shuffling of large genomic regions. Curr Opin Microbiol. 2017 Aug 1;38:22–9.

21. Matthey N, Stutzmann S, Stoudmann C, Guex N, Iseli C, Blokesch M. Neighbor predation linked to natural competence fosters the transfer of large genomic regions in Vibrio cholerae. Mignot T, Weigelp D, editors. eLife. 2019 Sep 3;8:e48212.

22. Godeux AS, Svedholm E, Barreto S, Potron A, Venner S, Charpentier X, et al. Interbacterial Transfer of Carbapenem Resistance and Large Antibiotic Resistance Islands by Natural Transformation in Pathogenic Acinetobacter. mBio. 2022 Jan 25;13(1):e02631–21.

23. Kloos J, Johnsen PJ, Harms K. Tn1 transposition in the course of natural transformation enables horizontal antibiotic resistance spread in Acinetobacter baylyi. Microbiology [Internet]. 2021 [cited 2024 Oct 10];167(1). Available from: https://www.ncbi.nlm.nih.gov/pmc/articles/PMC8116780/

24. Domingues S, Harms K, Fricke WF, Johnsen PJ, Silva GJ da, Nielsen KM. Natural Transformation Facilitates Transfer of Transposons, Integrons and Gene Cassettes between Bacterial Species. PLoS Pathog [Internet]. 2012 Aug [cited 2024 Oct 10];8(8). Available from: https://www.ncbi.nlm.nih.gov/pmc/articles/PMC3410848/

25. Kothari A, Soneja D, Tang A, Carlson HK, Deutschbauer AM, Mukhopadhyay A. Native Plasmid-Encoded Mercury Resistance Genes Are Functional and Demonstrate Natural Transformation in Environmental Bacterial Isolates. mSystems. 2019 Dec 17;4(6):10.1128/msystems.00588-19.

26. Maree M, Thi Nguyen LT, Ohniwa RL, Higashide M, Msadek T, Morikawa K. Natural transformation allows transfer of SCCmec-mediated methicillin resistance in Staphylococcus aureus biofilms. Nat Commun. 2022 May 5;13(1):2477.

27. Hülter N, Wackernagel W. Double illegitimate recombination events integrate DNA segments through two different mechanisms during natural transformation of Acinetobacter baylyi. Mol Microbiol. 2008;67(5):984–95.

28. Lloyd RG, Buckman C. Conjugational Recombination in Escherichia Coli: Genetic Analysis of Recombinant Formation in Hfr X F(-) Crosses. Genetics. 1995 Mar;139(3):1123.

29. Makky S, Dawoud A, Safwat A, Abdelsattar AS, Rezk N, El-Shibiny A. The bacteriophage decides own tracks: When they are with or against the bacteria. Curr Res Microb Sci. 2021 Dec 1;2:100050.

30. Dell’Annunziata F, Folliero V, Giugliano R, Filippis AD, Santarcangelo C, Izzo V, et al. Gene Transfer Potential of Outer Membrane Vesicles of Gram-Negative Bacteria. Int J Mol Sci. 2021 Jun 1;22(11):5985.

31. Denise R, Abby SS, Rocha EPC. Diversification of the type IV filament superfamily into machines for adhesion, protein secretion, DNA uptake, and motility. PLoS Biol [Internet]. 2019 Jul 19 [cited 2021 Feb 20];17(7). Available from: https://www.ncbi.nlm.nih.gov/pmc/articles/PMC6668835/

32. Carlson CA, Pierson LS, Rosen JJ, Ingraham JL. Pseudomonas stutzeri and related species undergo natural transformation. J Bacteriol. 1983 Jan;153(1):93–9.

33. Evans BA, Rozen DE. Significant variation in transformation frequency in Streptococcus pneumoniae. ISME J. 2013 Apr;7(4):791–9.

34. Sikorski J, Teschner N, Wackernagel W. Highly different levels of natural transformation are associated with genomic subgroups within a local population of Pseudomonas stutzeri from soil. Appl Environ Microbiol. 2002 Feb;68(2):865–73.

35. Maughan H, Redfield RJ. Extensive variation in natural competence in Haemophilus influenzae. Evol Int J Org Evol. 2009 Jul;63(7):1852–66.

36. Godeux AS, Lupo A, Haenni M, Guette-Marquet S, Wilharm G, Laaberki MH, et al. Fluorescence-Based Detection of Natural Transformation in Drug-Resistant Acinetobacter baumannii. J Bacteriol [Internet]. 2018 Sep 10 [cited 2021 Feb 18];200(19). Available from: https://www.ncbi.nlm.nih.gov/pmc/articles/PMC6148472/

37. Durieux I, Ginevra C, Attaiech L, Picq K, Juan PA, Jarraud S, et al. Diverse conjugative elements silence natural transformation in Legionella species. Proc Natl Acad Sci. 2019 Sep 10;116(37):18613–8.

38. Provvedi R, Chen I, Dubnau D. NucA is required for DNA cleavage during transformation of Bacillus subtilis. Mol Microbiol. 2001;40(3):634–44.

39. Nielsen KM, Johnsen PJ, Bensasson D, Daffonchio D. Release and persistence of extracellular DNA in the environment. Environ Biosafety Res. 2007 Jan;6(1–2):37–53.

40. Wintersdorff CJH von, Penders J, Niekerk JM van, Mills ND, Majumder S, Alphen LB van, et al. Dissemination of Antimicrobial Resistance in Microbial Ecosystems through Horizontal Gene Transfer. Front Microbiol. 2016 Feb 19;7:173.

41. Winter M, Buckling A, Harms K, Johnsen PJ, Vos M. Antimicrobial resistance acquisition via natural transformation: context is everything. Curr Opin Microbiol. 2021 Dec 1;64:133– 8.

42. Bi D, Xie R, Zheng J, Yang H, Zhu X, Ou HY, et al. Large-Scale Identification of AbaR-Type Genomic Islands in Acinetobacter baumannii Reveals Diverse Insertion Sites and Clonal Lineage-Specific Antimicrobial Resistance Gene Profiles. Antimicrob Agents Chemother. 2019 Mar 27;63(4):e02526–18.

43. Jorth P, Whiteley M. An Evolutionary Link between Natural Transformation and CRISPR Adaptive Immunity. mBio. 2012 Oct 2;3(5):10.1128/mbio.00309-12.

44. Croucher NJ, Harris SR, Barquist L, Parkhill J, Bentley SD. A High-Resolution View of Genome-Wide Pneumococcal Transformation. PLOS Pathog. 2012 Jun 14;8(6):e1002745.

45. Lin EA, Zhang XS, Levine SM, Gill SR, Falush D, Blaser MJ. Natural Transformation of Helicobacter pylori Involves the Integration of Short DNA Fragments Interrupted by Gaps of Variable Size. PLOS Pathog. 2009 Mar 13;5(3):e1000337.

46. Power JJ, Pinheiro F, Pompei S, Kovacova V, Yüksel M, Rathmann I, et al. Adaptive evolution of hybrid bacteria by horizontal gene transfer. Proc Natl Acad Sci U S A. 2021 Mar 9;118(10):e2007873118.

47. Johnston C, Martin B, Polard P, Claverys JP. Postreplication targeting of transformants by bacterial immune systems? Trends Microbiol. 2013 Oct 1;21(10):516–21.

48. Sánchez-Busó L, Comas I, Jorques G, González-Candelas F. Recombination drives genome evolution in outbreak-related Legionella pneumophila isolates. Nat Genet. 2014 Nov;46(11):1205–11.

49. David S, Rusniok C, Mentasti M, Gomez-Valero L, Harris SR, Lechat P, et al. Multiple major disease-associated clones of Legionella pneumophila have emerged recently and independently. Genome Res. 2016 Nov 1;26(11):1555–64.

50. Snitkin ES, Zelazny AM, Montero CI, Stock F, Mijares L, NISC Comparative Sequence Program, et al. Genome-wide recombination drives diversification of epidemic strains of Acinetobacter baumannii. Proc Natl Acad Sci. 2011 Aug 16;108(33):13758–63.

51. Johnston C, Martin B, Granadel C, Polard P, Claverys JP. Programmed Protection of Foreign DNA from Restriction Allows Pathogenicity Island Exchange during Pneumococcal Transformation. PLOS Pathog. 2013 Feb 14;9(2):e1003178.

52. Blokesch M, Schoolnik GK. Serogroup Conversion of Vibrio cholerae in Aquatic Reservoirs. PLOS Pathog. 2007 Jun 8;3(6):e81.

53. Miller MC, Keymer DP, Avelar A, Boehm AB, Schoolnik GK. Detection and transformation of genome segments that differ within a coastal population of Vibrio cholerae strains. Appl Environ Microbiol. 2007;73(11):3695–704.

54. Vorwerk H, Huber C, Mohr J, Bunk B, Bhuju S, Wensel O, et al. A transferable plasticity region in ampylobacter coli allows isolates of an otherwise non-glycolytic food-borne pathogen to catabolize glucose. Mol Microbiol. 2015;98(5):809–30.

55. Lu J, Wang Y, Zhang S, Bond P, Yuan Z, Guo J. Triclosan at environmental concentrations can enhance the spread of extracellular antibiotic resistance genes through transformation. Sci Total Environ. 2020 Apr 15;713:136621.

56. Leidy G, Hahn E, Alexander HE. ON THE SPECIFICITY OF THE DESOXYRIBONUCLEIC ACID WHICH INDUCES STREPTOMYCIN RESISTANCE IN HEMOPHILUS. J Exp Med. 1956 Sep 1;104(3):305.

57. Janoir C, Podglajen I, Kitzis MD, Poyart C, Gutmann L. In Vitro Exchange of Fluoroquinolone Resistance Determinants between Streptococcus pneumoniae and Viridans Streptococci and Genomic Organization of the parE-parC Region in S. mitis. J Infect Dis. 1999 Aug 1;180(2):555–8.

58. Bowler LD, Zhang QY, Riou JY, Spratt BG. Interspecies recombination between the penA genes of Neisseria meningitidis and commensal Neisseria species during the emergence of penicillin resistance in N. meningitidis: natural events and laboratory simulation. J Bacteriol. 1994 Jan;176(2):333–7.

59. Traglia GM, Place K, Dotto C, Fernandez JS, Montaña S, Bahiense C dos S, et al. Interspecies DNA acquisition by a naturally competent Acinetobacter baumannii strain. Int J Antimicrob Agents. 2019 Jan 3;53(4):483.

60. Perez F, Stiefel U. The Impact of Natural Transformation on the Acquisition of Antibiotic Resistance Determinants. mBio. 2022 May 12;13(3):e00336–22.

61. Seemann T. Prokka: rapid prokaryotic genome annotation. Bioinforma Oxf Engl. 2014 Jul 15;30(14):2068–9.

62. Perrin A, Rocha EPC. PanACoTA: a modular tool for massive microbial comparative genomics. NAR Genomics Bioinforma [Internet]. 2021 Feb 1 [cited 2021 Feb 17];3(lqaa106). Available from: 10.1093/nargab/lqaa106

63. Croucher NJ, Page AJ, Connor TR, Delaney AJ, Keane JA, Bentley SD, et al. Rapid phylogenetic analysis of large samples of recombinant bacterial whole genome sequences using Gubbins. Nucleic Acids Res. 2015 Feb 18;43(3):e15.

64. Mostowy R, Croucher NJ, Andam CP, Corander J, Hanage WP, Marttinen P. Efficient Inference of Recent and Ancestral Recombination within Bacterial Populations. Mol Biol Evol. 2017 May;34(5):1167–82.

65. Ishikawa SA, Zhukova A, Iwasaki W, Gascuel O. A Fast Likelihood Method to Reconstruct and Visualize Ancestral Scenarios. Mol Biol Evol. 2019 Sep 1;36(9):2069–85.

66. Neron B, Denise R, Coluzzi C, Touchon M, Rocha EPC, Abby SS. MacSyFinder v2: Improved modelling and search engine to identify molecular systems in genomes [Internet]. bioRxiv; 2023 [cited 2023 Mar 1]. p. 2022.09.02.506364. Available from: https://www.biorxiv.org/content/10.1101/2022.09.02.506364v3

67. Abby SS, Néron B, Ménager H, Touchon M, Rocha EPC. MacSyFinder: A Program to Mine Genomes for Molecular Systems with an Application to CRISPR-Cas Systems. PLOS ONE. 2014 Oct 17;9(10):e110726.

68. Guo J, Bolduc B, Zayed AA, Varsani A, Dominguez-Huerta G, Delmont TO, et al. VirSorter2: a multi-classifier, expert-guided approach to detect diverse DNA and RNA viruses. Microbiome. 2021 Feb 1;9(1):37.

69. Nayfach S, Camargo AP, Schulz F, Eloe-Fadrosh E, Roux S, Kyrpides NC. CheckV assesses the quality and completeness of metagenome-assembled viral genomes. Nat Biotechnol. 2021 May;39(5):578–85.

70. Xie Z, Tang H. ISEScan: automated identification of insertion sequence elements in prokaryotic genomes. Bioinformatics. 2017 Nov 1;33(21):3340–7.

71. Feldgarden M, Brover V, Gonzalez-Escalona N, Frye JG, Haendiges J, Haft DH, et al. AMRFinderPlus and the Reference Gene Catalog facilitate examination of the genomic links among antimicrobial resistance, stress response, and virulence. Sci Rep. 2021 Jun 16;11(1):12728.

72. Tung Ho L si, Ané C. A Linear-Time Algorithm for Gaussian and Non-Gaussian Trait Evolution Models. Syst Biol. 2014 May 1;63(3):397–408.

