## Supplemental Figures for "Impact of natural transformation on the acquisition of novel genes in bacteria"

**List of Supplementary Tables**

**Table S1** Bacterial strains and their phenotypic and genomic repertoire features.

**Table S2** Association of transformation phenotype with genomic repertoire features in *Acinetobacter baumannii* and *Legionella pneumophila* with phylolm.

**Table S3** Counts of the number of accessory genes found in each genomic context depending on the phenotype in *Acinetobacter baumannii* and *Legionella pneumophila*.

**List of Supplementary Figures**

**Figure S1.** Recombination contexts of accessory genes that can be observed depending on the draft assemblies.

**Figure S2.** Scheme depicting the estimation of transformation contribution to gene gains in *Legionella pneumophila* and *Acinetobacter baumannii*.

**Figure S3.** Number of acquisition events whose gene gains are interrupted or not by genes not classified as gains in *Acinetobacter baumannii* and *Legionella pneumophila*.

**Figure S4.** Hotspots of gene gains in transformable and non-transformable strains in *Acinetobacter baumannii* and *Legionella pneumophila* along their respective reference genomes.

**Figure S5.** Characterization of hotspot of gene gains.

**Figure S6.** Distribution of the number of conjugative systems per genome and characterization of their types *in Acinetobacter baumannii* and *Legionella pneumophila*.

**Figure S7.** Distribution of the number of prophages per genome in *Acinetobacter baumannii*.

**Figure S8.** Distribution of the number of accessory genes that are not gains in between consecutive gene gains within intervals in *Acinetobacter baumannii* (left) and *Legionella pneumophila* (right).

**Figure S9.** Distribution of the number of genes in insertion sequences interrupting acquisition events whose genes were not marked as gains in *Acinetobacter baumannii* and *Legionella pneumophila*.

**Figure S10.** Distribution of the number of clusters of gains defined in intervals presenting gene gains in *Acinetobacter baumannii* (left) and *Legionella pneumophila* (right).

**Figure S11.** Definition of hotspots and coldspots.

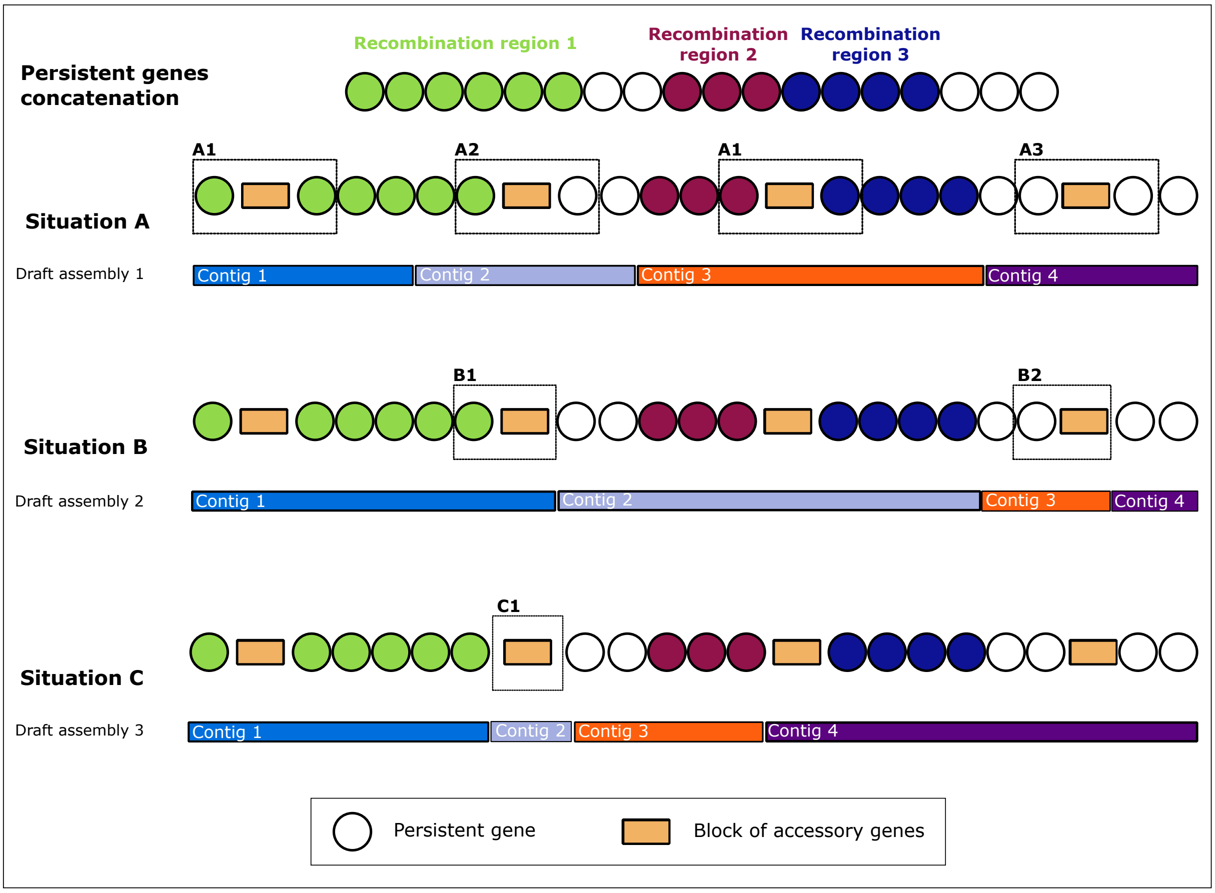


**Figure S1. Recombination contexts of accessory genes that can be observed depending on the draft assemblies.** Persistent genes are represented as circles, colored when part of a recombination tract and white when they are not. Orange rectangles stand for blocks of accessory genes.

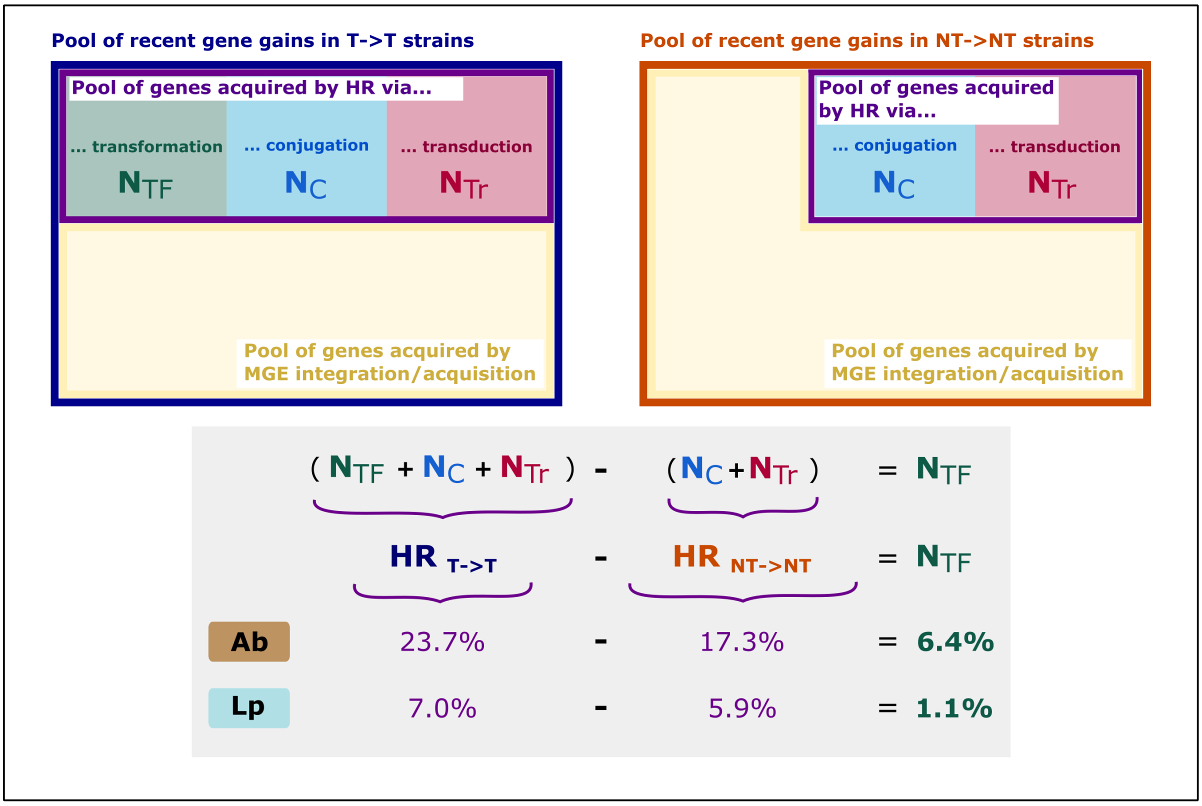


**Figure S2. Scheme depicting the estimation of transformation contribution to gene gains in *Legionella pneumophila* and *Acinetobacter baumannii*.** HR stands for homologous recombination. $N_{TF}$, $N_{C}$ and $N_{Tr}$ are the number of gains acquired by transformation, conjugation and transduction respectively.

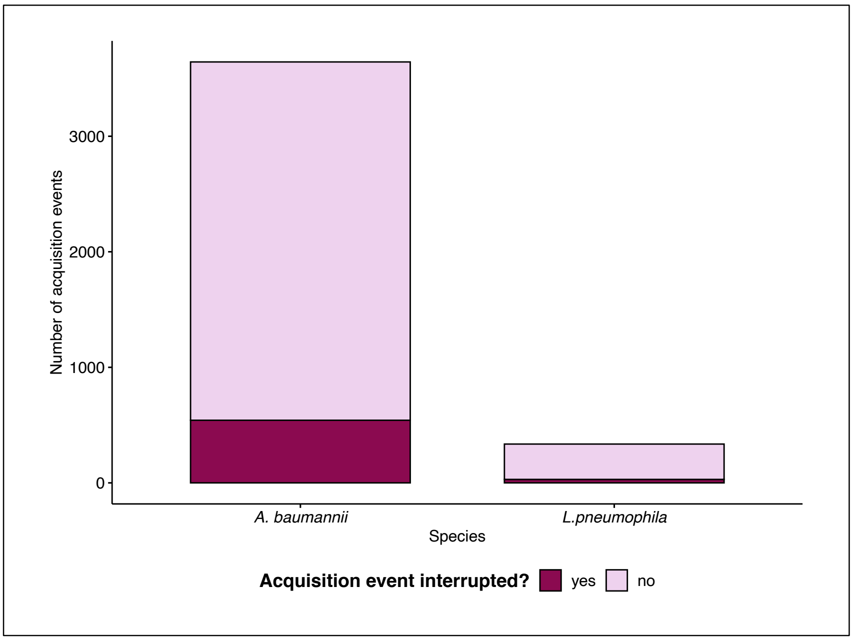


**Figure S3. Number of acquisition events whose gene gains are interrupted or not by genes not classified as gains in *Acinetobacter baumannii* and *Legionella pneumophila.***

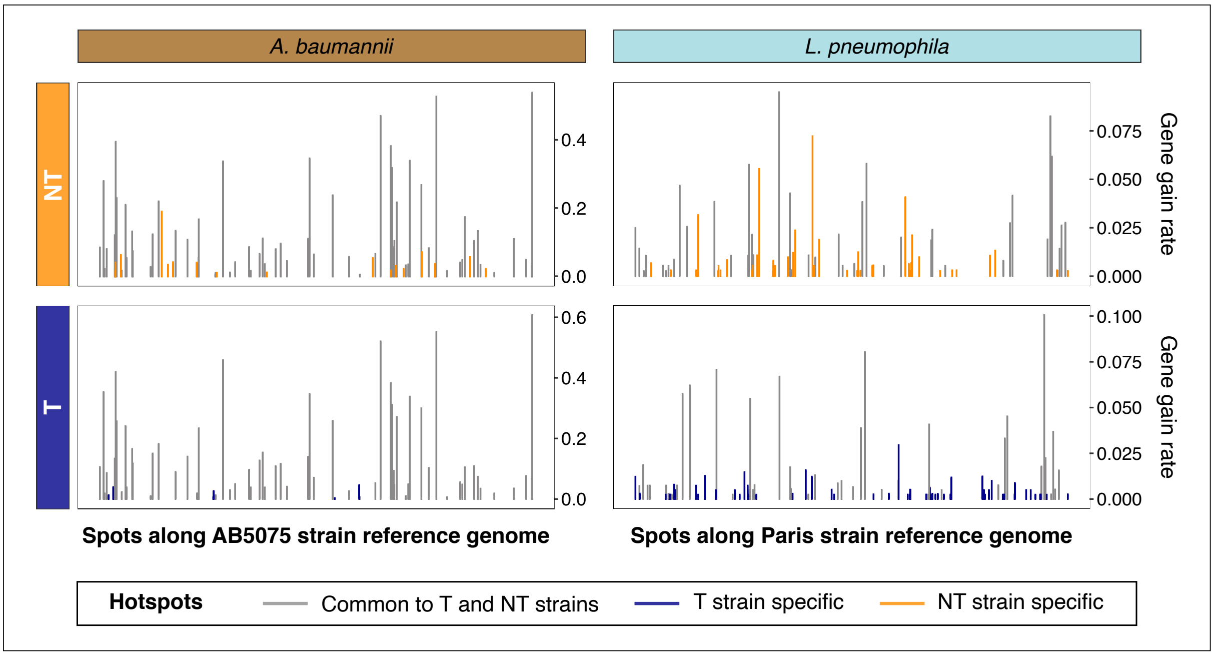


**Figure S4. Hotspots of gene gains in transformable and non-transformable strains in *Acinetobacter baumannii* and *Legionella pneumophila* along their respective reference genomes.**

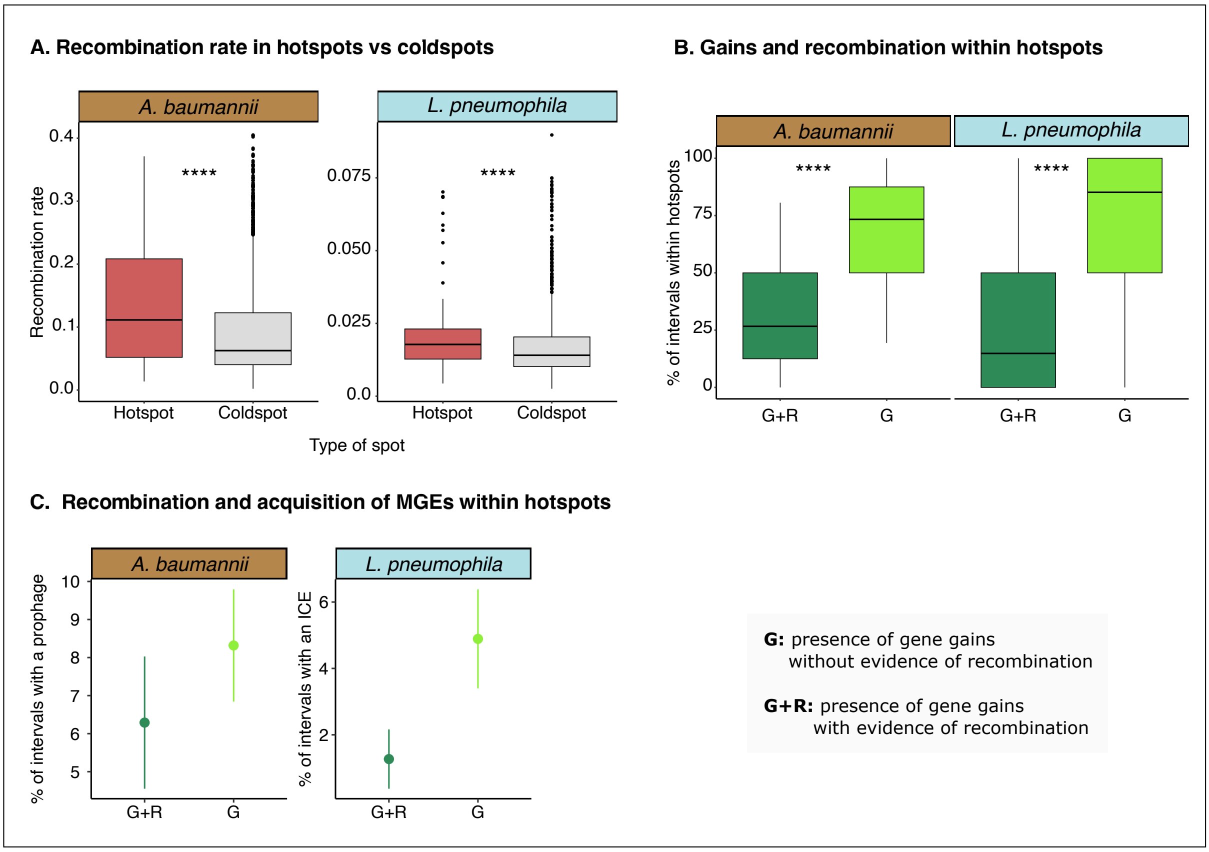


**Figure S5. Characterization of hotspot of gene gains.**
**A. Distribution of recombination rates in persistent genes flanking hotspots and coldspots defined on all genomes.**
**B. Comparison of the proportion of intervals with at least one gene gain in hotspots of gains (defined on all genomes) that present evidence of recombination and the ones that do not.**
**C. Proportion of intervals with gains within hotspots (defined on all genomes) that are carrying recently acquired large MGEs (prophage in Ab and ICE in Lp) depending on the interval presenting evidence of recombination or not.** Dots stands for the average proportions and bars for the standard error. Statistical tests are Wilcoxon tests with ***: p<0.001, **: p<0.01**.**

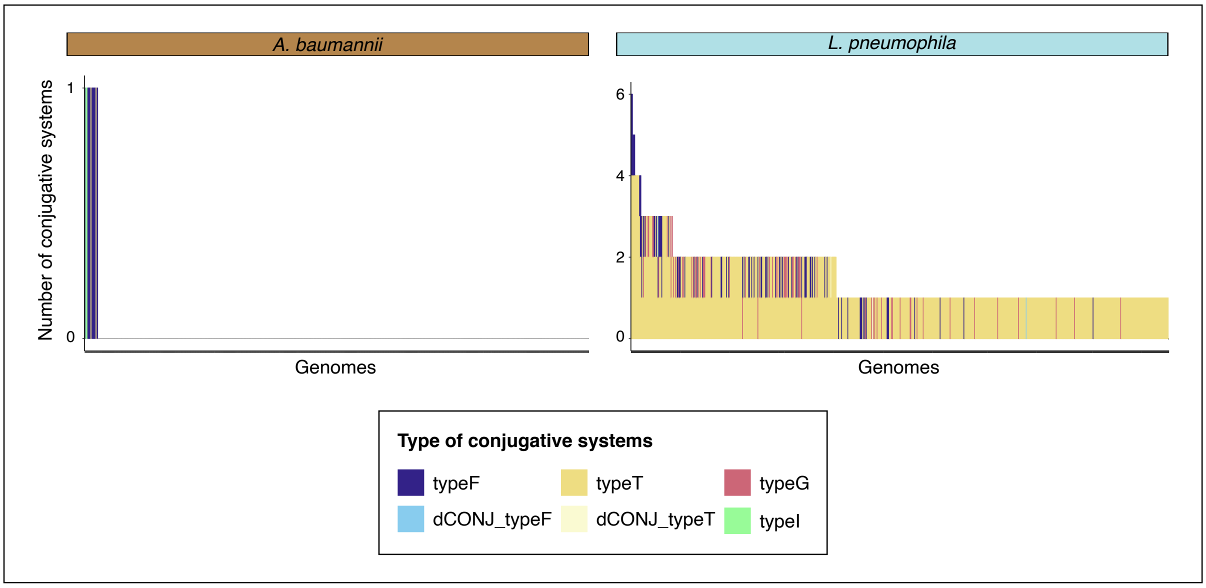


**Figure S6. Distribution of the number of conjugative systems per genome and characterization of their types in *Acinetobacter baumannii* and *Legionella pneumophila.***

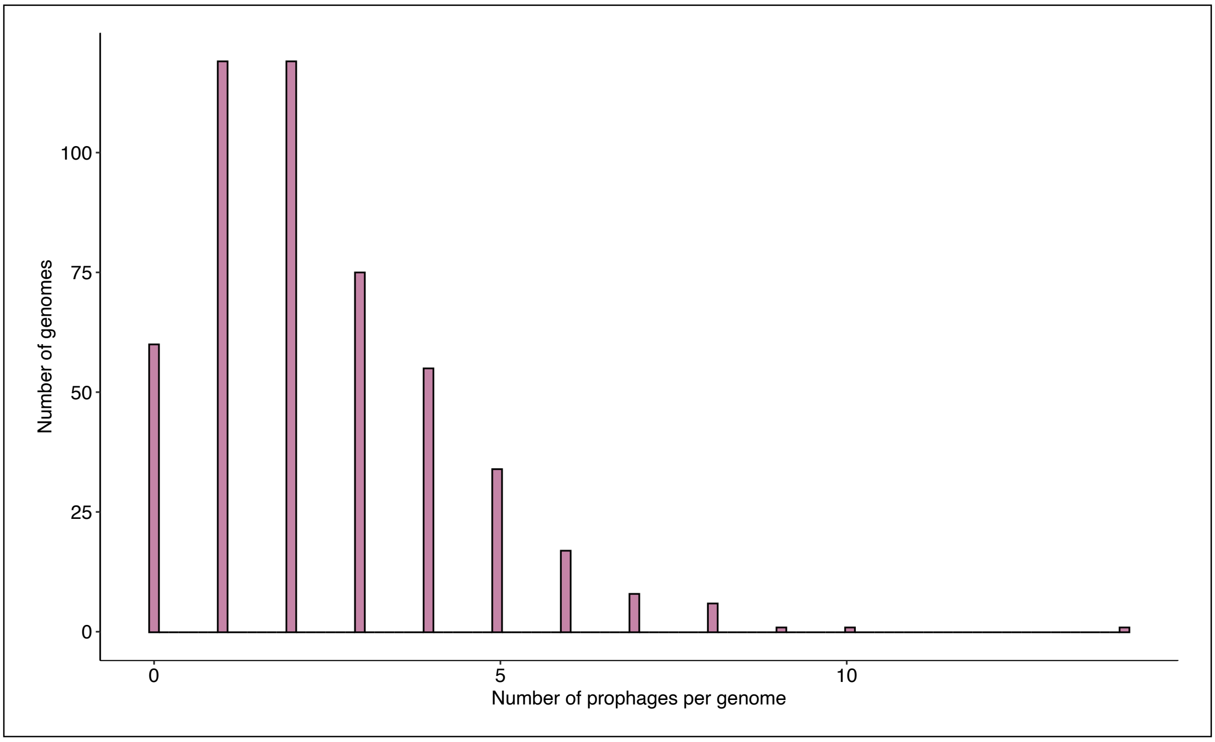


**Figure S7. Distribution of the number of prophages per genome in *Acinetobacter baumannii.***

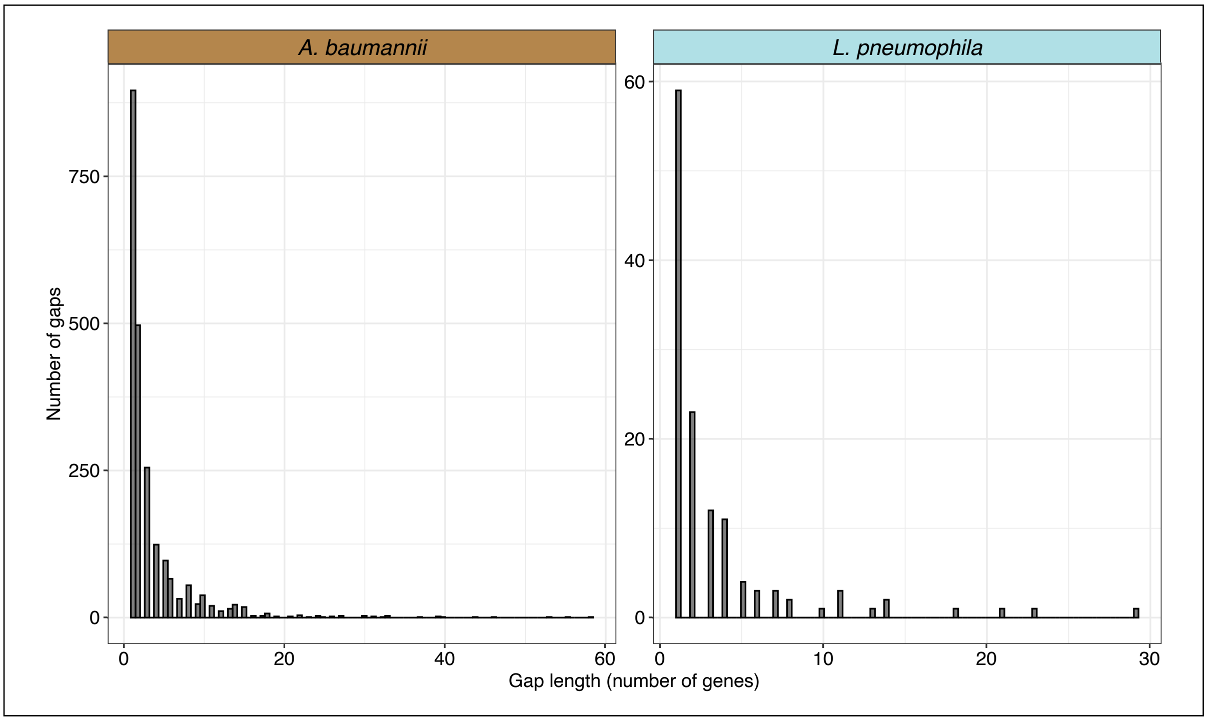


**Figure S8. Distribution of the number of accessory genes that are not gains in between consecutive gene gains within intervals in *Acinetobacter baumannii* (left) and *Legionella pneumophila* (right).**

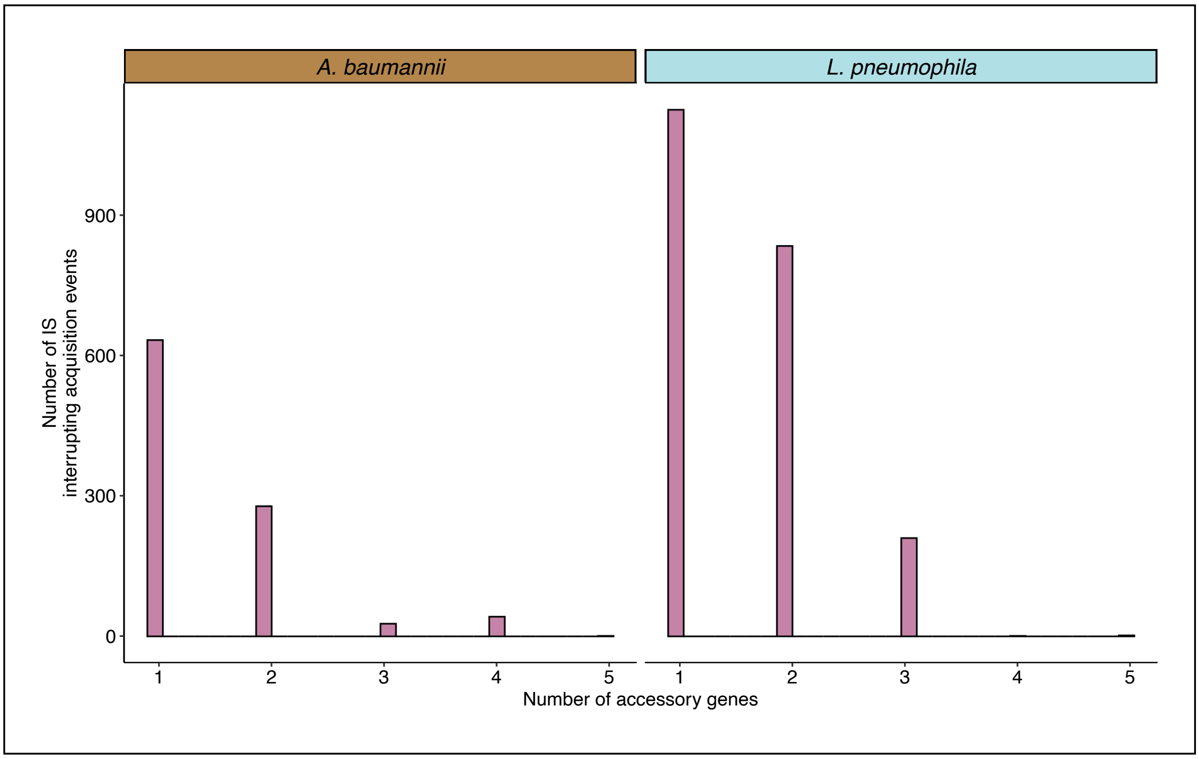


**Figure S9. Distribution of the number of genes in insertion sequences interrupting acquisition events whose genes were not marked as gains in *Acinetobacter baumannii* and *Legionella pneumophila*.**

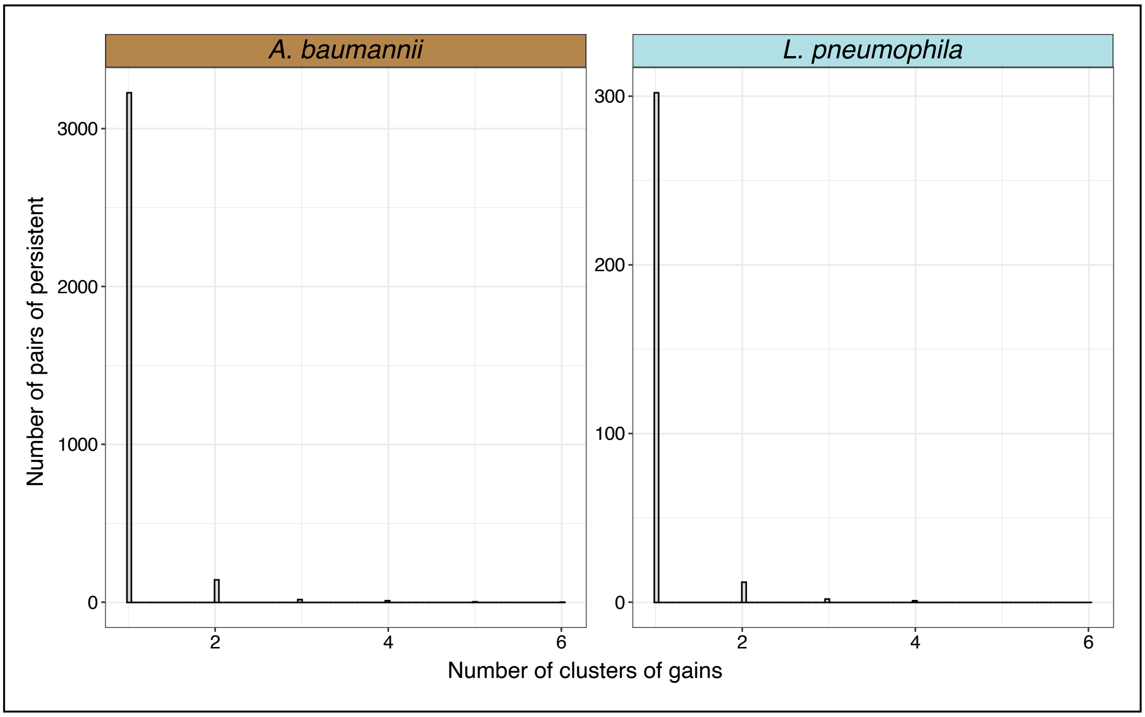


**Figure S10. Distribution of the number of clusters of gains defined in intervals presenting gene gains in *Acinetobacter baumannii* (left) and *Legionella pneumophila* (right).**

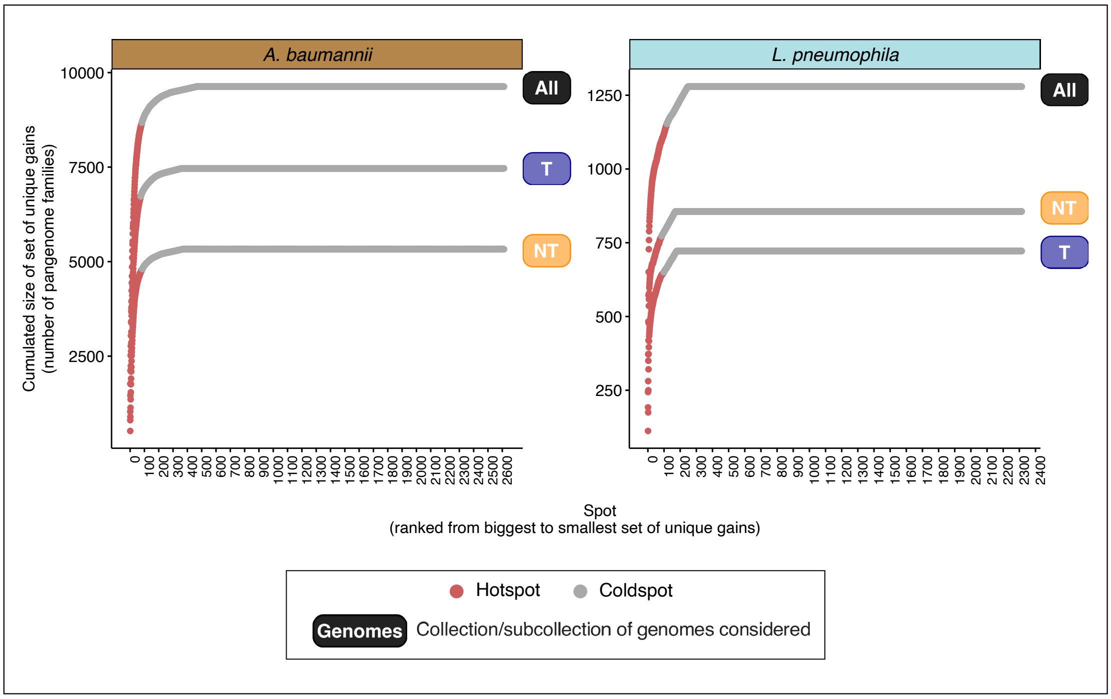


**Figure S11. Definition of hotspots and coldspots.** Definitions are based on the cumulative distribution function of the numbers of unique gained pangenome families in spots. Spots were identified in *Acinetobacter baumannii* and *Legionella pneumophila* collections and subcollections. Spots considered as hotspots were colored in red while the coldspots were in grey. The cumulated size of the set of unique gains in spots was computed for the spots observed in all genomes (All), the ones observed in the genomes of transformable strains (T) and the ones observed in the genomes of non-transformable strains (NT) and all the three cumulated curves were plotted on the same graph for each species.
